# Effect of Photoinitiation Process on Photo-Crosslinking of Gelatin Methacryloyl Hydrogel Networks

**DOI:** 10.1101/2025.04.10.648118

**Authors:** Doğukan Duymaz, İsmail Can Karaoğlu, Seda Kizilel

## Abstract

Gelatin methacryloyl (GelMA) has emerged as a widely utilized biomaterial in tissue engineering due to its tunable mechanical properties, cell-adhesive motifs, and photo-crosslinkability. However, the physicochemical characteristics and biomedical utility of GelMA hydrogels are greatly influenced by the choice and concentration of photoinitiating systems. Despite the increasing acceptance of visible-light and UV-sensitive initiators, a systematic comparative evaluation of their impact on GelMA hydrogel properties remains limited. In this study, we present the first systematic investigation of how individual photoinitiators, Eosin Y (EY), Lithium Phenyl-2,4,6-trimethylbenzoylphosphinate (LAP), Ruthenium (II) trisbipyridyl chloride ([RuII(bpy)_3_]^2+^) (Ru), affect the viscoelastic properties, swelling behavior, degradation kinetics, and cytocompatibility of 5% and 10% (w/v) GelMA hydrogels. By varying photoinitiator concentrations ([EY]: 0.005–0.1 mM, [LAP]: 0.01–0.5% (w/v), [Ru]: 0.02–1 mM) and utilizing consistent light intensity (10 mW/cm^2^ at system-specific wavelengths), we identified critical thresholds and plateau behaviors that distinctly influenced the stiffness and integrity of the hydrogels. Our findings revealed that each photoinitiating system exhibits unique advantages and trade-offs. LAP and Ru systems facilitated rapid gelation with easier utilization and were associated with higher swelling and accelerated degradation profiles—features particularly advantageous for applications such as 3D bioprinting and *in situ* injectable hydrogel systems. However, their atypical behaviors at certain concentrations and light exposure durations highlight the necessity for precise control and further mechanistic exploration. In contrast, EY-mediated hydrogels offered superior stiffness and minimal swelling at optimal concentrations, favoring applications that demand long-term mechanical stability, at the cost of a more complex cross-linking mechanism. Notably, by correlating mechanical and degradation behaviors with NIH-3T3 fibroblast viability, we also assessed biocompatibility window for each concentration of the systems, linking biomaterial performance with biomedical applicability. Overall, our study underlines the importance of tailoring photoinitiator selection and concentration to specific application needs, striking a balance between gelation kinetics, mechanical integrity, degradation behavior, and cytocompatibility. These insights provide a foundational framework for engineering GelMA-based hydrogels paving the way for reproducible, efficient, targeted biomedical applications.

**Graphical Abstract:** 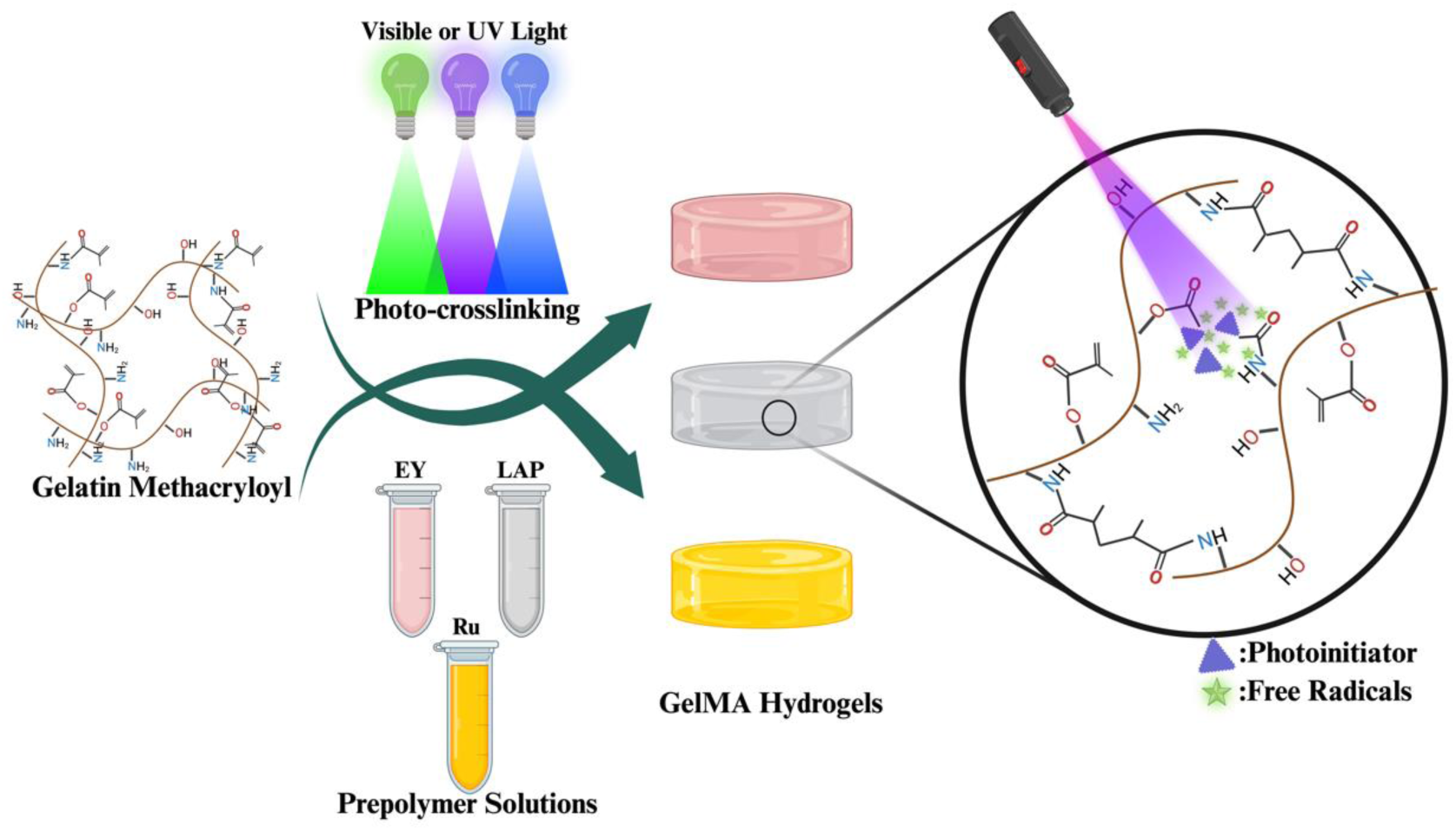

## 1. Introduction

Hydrogels are 3D crosslinked polymer networks that absorb and retain large amounts of water due to their hydrophilic groups (-NH2, -COOH, -OH, -CONH2, -CONH, etc.). Their high-water content, porosity, and tissue-like structure resulting from chemical or physical cross-linking of natural or synthetic polymers make them soft and flexible.^1^ Over the past 60 years, numerous types of hydrogels have emerged as critical platforms for biomedical applications, especially tissue engineering, due to their tunable mechanical properties, biocompatibility, and ability to mimic the extracellular matrix (ECM) of biological tissues. ^1, 2^

Gelatin, a biodegradable and FDA-approved natural polymer derived from the partial degradation of collagen, is an exceptionally appealing choice for hydrogel fabrication. Its numerous advantages—including abundant availability, safety, high RGD content (Arg-Gly-Asp) and ease of procurement—make it a standout among protein-based polymers. ^3^ Furthermore, gelatin features a wealth of advantageous functional groups in its structure, including terminal -NH2 and -COOH groups, as well as aspartic acid (-COOH), lysine (-NH2), histidine (imidazolium), and arginine (guanidinium). This rich array of functional groups allows for a virtually limitless range of conjugation and modification strategies, enhancing the versatility and functionality of gelatin in various applications. ^3^ However, despite many advantages, pure gelatin hydrogels suffer from intrinsic limitations—such as insufficient mechanical strength and instability at body temperature—which significantly restricts its further biomedical applicability. ^4^ To address these drawbacks, an efficient modification replacing amine and hydroxyl groups of side chains of gelatin with methacrylic anhydride (MA), was presented by van den Bulcke *et al.* to obtain a photo-crosslinkable form of gelatin, termed later in literature as gelatin methacryloyl (GelMA). ^5^ Since then, GelMA has emerged as one of the most extensively studied biomaterials in tissue engineering, with continuous optimizations aimed at enhancing its robustness and expanding its range of applications. ^6, 7, 8, 9, 10^

The introduction of methacryloyl substituent groups grants gelatin a photo-crosslinking capability, enabling the formation of a stable 3D hydrogel network through free radical photopolymerization. The process is initiated by a photoinitiator or co-initiator activated by UV or visible light, which then generates free radicals to drive the polymerization of monomers or macromers, resulting in a crosslinked structure. ^5, 11^ Key photopolymerization parameters, including photoinitiator type and concentration, as well as light intensity and exposure time, are known to significantly influence the properties of the resulting hydrogels. This high degree of tunability enables the fabrication of GelMA hydrogels with tailored characteristics, making them highly versatile for a wide range of tissue engineering applications (**Figure 1a**). ^12, 13, 14^

**Figure 1.**
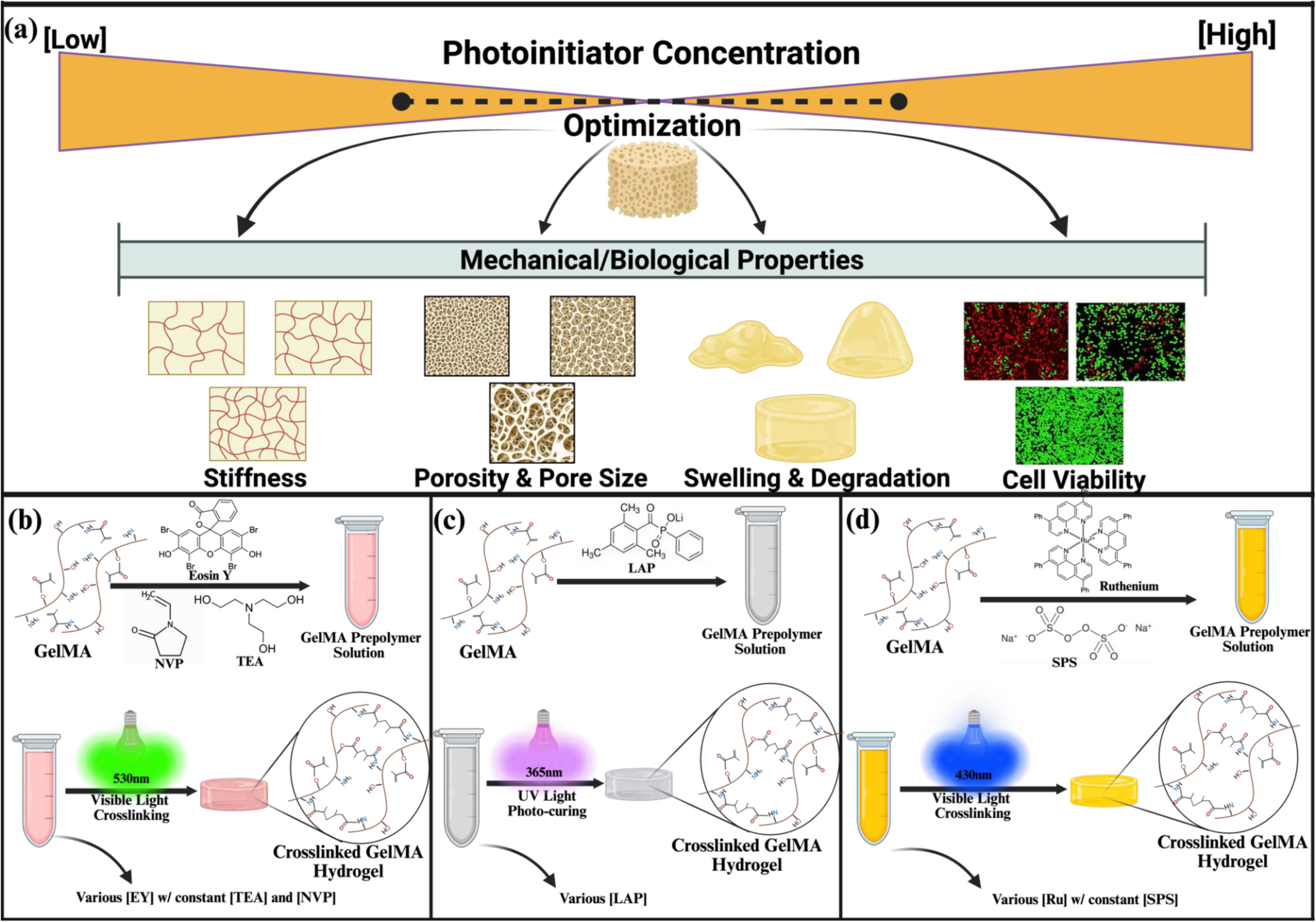
Schematic illustration of the influence of photoinitiator type and concentration on the mechanical and biological properties of GelMA hydrogels. (**a**) Optimization of photoinitiator concentration is critical for achieving balanced hydrogel properties, including mechanical stiffness, pore architecture (porosity and pore size), swelling behavior, enzymatic degradation resistance, and cytocompatibility. A bell-shaped concentration profile highlights the existence of an optimal range, beyond which excessive or insufficient radical formation leads to structural and biological instability. (**b**) EY photoinitiating system, employing NVP and TEA, activated under 530 nm green light. (**c**) LAP-mediated system crosslinked under 365 nm UV light. (**d**) Ru/SPS photoinitiating system initiated under 430 nm blue light.

Over the past two decades, various photoinitating systems have been utilized for the effective photo-crosslinking of acrylated materials like GelMA and PEGDA, with most relying on UV-light-induced activation. ^13, 15^ However, UV light exposure poses potential risks to both delivered cells and host tissues, as both prolonged long wavelength (UVA, 320-400nm) or short wavelength (UVB, <320 nm) irradiation can generate free radicals that induce DNA damage and impair cellular function. ^16, 17^ For instance, although Irgacure 2959 (2-hydroxy-4′- (2-hydroxyethoxy)-2-methylpropiophenone) (I2959) still remains as one of the most widely used photoinitiators in tissue engineering ^18^, its peak absorption around 280 nm and the resulting need of prolonged UVA exposure for effective cross-linking can induce cytotoxic effects, necessitating safer GelMA photopolymerization strategies in research recently. ^19, 20, 21, 22^ In this regard, the use of visible light photoinitiators such as Eosin Y (EY) ^23^, Lithium phenyl-2,4,6-trimethylbenzoylphosphinate (LAP) ^24^, and Ruthenium (II) trisbipyridyl chloride ([RuII(bpy)_3_]^2+^) (Ru) ^25^ has emerged as promising alternatives for fabrication of GelMA hydrogels efficiently. Moreover, visible or longer wavelength of UVA light offer superior tissue penetration and operates at lower energy levels compared to UVB light, making it particularly advantageous for *in situ* hydrogel formation and minimally invasive biomedical applications ^26^.

EY, an FDA-approved dye, facilitates photopolymerization through Type II photoinitiation mechanism when paired with a co-initiator triethanolamine (TEA). ^19^ Upon green light excitation (450-550 nm), EY facilitates the generation of TEA radicals, which drive branching and cross-linking reactions in the prepolymer, leading to gel formation. ^23, 27^ To enhance the gelation kinetics of the process, N-vinyl-2-pyrrolidone (NVP) or N-Vinyl caprolactam (NVC) are commonly introduced as a comonomer, increasing the concentration of vinyl groups in the reaction medium and synergistically cross-propagating with GelMA methacrylate groups (**Figure 1b**). ^28^ LAP, on the other hand, has an absorption peak around 365 nm, allowing it to be activated by low-intensity UV or blue light. ^29^ Numerous studies have highlighted the enhanced efficiency and reduced cytotoxicity of LAP-mediated photopolymerization of hydrogels compared to the widely used I2959. This superiority is attributed to LAP’s higher water solubility and greater molar extinction coefficient at 365 nm, facilitating more efficient cross-linking under UVA or even 405 nm blue light exposure. ^20, 30, 31^ Furthermore, LAP operates through a rapid Type I photoinitiation mechanism upon exposure to UVA or visible light, eliminating the need for a co-initiator or accelerator monomer, thereby enhancing its practicality and efficiency in photopolymerization (**Figure 1c**). ^31^ Finally, the Ru-based photoinitiation system has gained increasing prominence for the efficient cross-linking of GelMA and other biomaterials, driven by its expanding use in tissue engineering applications, including the fabrication of cell-laden scaffolds and bioinks for 3D bioprinting. ^25, 32, 33^ This process involves incorporating Ru as the photoinitiator alongside a persulfate-based co-initiator, such as ammonium or sodium persulfate (SPS), into polymers containing phenolic residues (e.g., tyrosine) and/or acrylated groups, followed by exposure to visible light within the 400–500 nm range (**Figure 1d**). ^34, 35^ Building on previous promising studies that have demonstrated the Ru system’s efficiency in cross-linking protein-based biopolymers ^36, 37^, we have recently utilized this strategy to develop a novel clinical treatment of keratoconus through the photopolymerization of collagen layer in cornea with Ru-based visible light treatment instead of using cytotoxic UV exposure. ^38, 39^ Additionally, Lim *et al.* revealed the fast kinetics and efficiency of Ru-based photopolymerization on GelMA hydrogels, outperforming I2959 and LAP-mediated photoinitiator systems. ^26^ Their studies later underline the necessity of further systematic research into the underlying mechanisms and the resulting mechanical properties of hydrogels crosslinked using the Ru/SPS system.

Despite their similar mechanism in free radical photopolymerization, the type and concentration of these photoinitiators and its components are known to significantly influence resulting gel properties such as mechanical strength, degradation behavior, printability, or cell compatibility. ^12,35^ However, a comparative analysis of individual photoinitiators, focusing on key hydrogel characteristics—including viscoelasticity, swelling behavior, biodegradation profile, and cytocompatibility—has yet to be performed to carefully tailor to the specific biomedical application. Previously, we investigated the impact of individual concentrations of key parameters within the EY photoinitiating system on GelMA hydrogels to train an Artificial Neural Network (ANN) model, which functions as a comprehensive tool for predicting the elastic modulus and gelation time of the resulting hydrogels. ^40^ This study provided critical understandings into how photopolymerization efficiency and consequently, the final properties of the hydrogel are strongly dictated by both the type and concentration of the photoinitiator and its components.

Building upon this understanding, the present study aims to systematically investigate the effects of three distinct photoinitiators (EY, LAP, and Ru), on key characteristics of 5% and 10% (w/v) GelMA hydrogels, including real-time stiffness and gelation kinetics, degradation rate, swelling behavior, and cytocompatibility. By altering photoinitiator concentrations ([EY]: 0.005–0.1 mM, [LAP]: 0.01–0.5% (w/v), [Ru]: 0.02–1 mM), for the first time in the literature, we provide a comprehensive comparative analysis of these photoinitiators together, within the same biomaterial system cross-linked via free radical photopolymerization.

Based on our previous findings and literature, we selected the most cytocompatible concentrations for both the photoinitiators and the associated components of each photoinitiating system, specifically 100 mM TEA and 50 mM NVP for the EY system, and a constant 1:10 molar ratio of Ru:SPS across all tested concentrations. We observed distinct gelation profiles for 5% and 10% (w/v) GelMA hydrogels across EY-, LAP-, and Ru-based photoinitiating systems, each exhibiting unique advantages and trade-offs about characteristics of fabricated hydrogels. We validated our previous findings that EY concentrations enhance stiffness up to a critical threshold (0.02 mM), beyond which stiffness declines. ^40^ Similarly, 0.5 mM was identified as the critical upper limit for Ru, above which mechanical strength significantly diminished. In the LAP system, stiffness increased proportionally with concentration under short UV exposure, whereas prolonged exposure led to atypical gelation behavior, particularly in lower concentrations (0.01, and 0.02% (w/v)). These mechanical profiles were validated by enzymatic degradation and swelling analyses, where stiffer gels demonstrated greater resistance to enzymatic breakdown and lower swelling ratios. To assess biocompatibility, NIH-3T3 mouse fibroblast cells were seeded on hydrogels prepared under each condition, enabling the identification of both optimal and cytotoxic concentration ranges.

In summary, this study is particularly significant as it offers critical insights into the rational design and optimization of GelMA hydrogels across various photoinitiation systems. By elucidating the distinct influences of different photoinitiators, our findings will serve as a valuable reference for researchers in the selection and fine-tuning of photoinitiators for GelMA cross-linking, ultimately enabling more reproducible, efficient, and clinically-translatable advancements in tissue engineering, bioprinting, and regenerative medicine.

## 2. Results and Discussion

### 2.1. GelMA Characterization

The successful methacrylation of gelatin was characterized through ^1^H NMR, FT-IR and XRD analyses as presented in **Figure 2**. The appearance of distinct peaks corresponding to the methacrylate vinyl protons is observed at δ = 5.4 and 5.6 ppm (yellow, denoted as (**1**)) and δ = 1.9 ppm (purple, denoted as (**3**)), indicating the incorporation of methacrylate functional groups (**Figure 2a**). Additionally, a decrease in the signal intensity at δ = 2.9 ppm (green, denoted as (**2**)), which corresponds to the methylene protons of lysine residues in gelatin, further supports the modification. By comparing the signal at δ = 2.9 ppm between unmodified gelatin and GelMA, the degree of methacrylation was approximately quantified as 81.4 ± 1.2%, demonstrating a high extent of functionalization.

**Figure 2.**
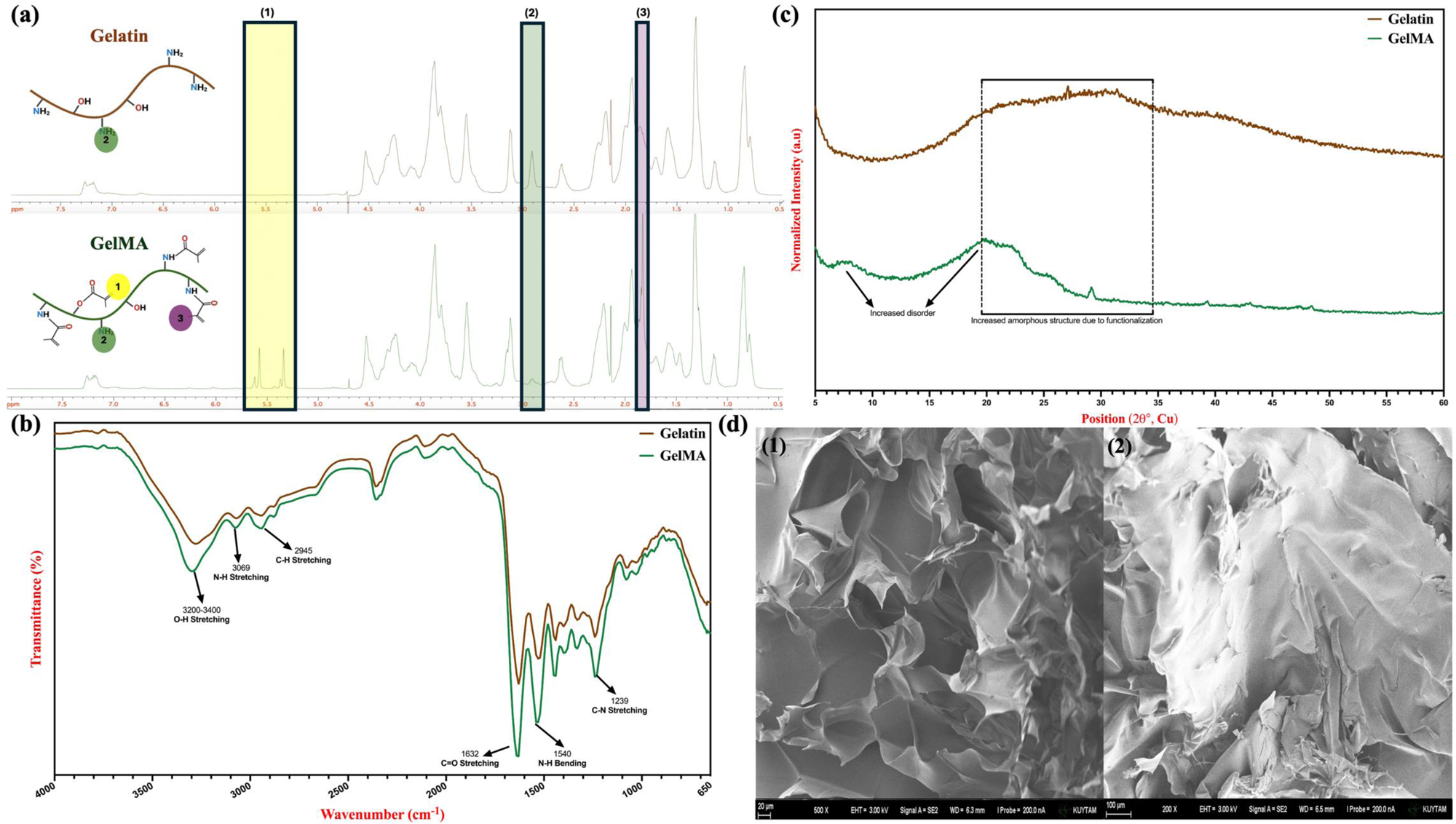
Synthesized GelMA characterization. (**a**) H-NMR spectrum of GelMA, highlighting the characteristic protons of the methacrylate vinyl group at δ = 1.9 ppm (3, purple), 5.4 ppm, and 5.6 ppm (1, yellow), along with the methylene protons of lysine at δ = 2.9 ppm (2, green). (**b**) FTIR spectrum of GelMA, displaying key absorption peaks at 3290 cm^-1^ (O–H stretching vibration), 3200–3400 cm^-1^ (N–H stretching), 3069 cm^-1^ (C–H stretching vibration), 1640 cm^-1^ (C=O stretching), 1541 cm^-1^ (N–H bending), and 1240 cm^-1^ (C–N stretching). (**c**) XRD pattern of gelatin and GelMA demonstrating crystallinity difference of the natural and functionalized polymer. (**d**) FE-SEM images of GelMA, showing (1) the cross-sectional structure and (2) the top-view morphology.

In the FT-IR spectra of gelatin and GelMA, characteristic absorption bands corresponding to the chemical structure of both materials were observed (**Figure 2b**). The broad peak observed at 3200–3400 cm⁻¹, corresponding to O-H stretching from hydroxyproline units, along with the band at approximately 3069 cm⁻¹, attributed to N-H stretching of peptide bonds (amide A), appeared in both the gelatin and GelMA spectra. However, a noticeable shift was observed in the GelMA spectrum, indicating modifications resulting from methacrylation. Additionally, the similar shift peak at 2945 cm⁻¹, attributed to the symmetric and asymmetric stretching of CH₂ groups in alkyl chains, further supports the structural modifications introduced by methacrylation. The GelMA spectrum also exhibits distinct peaks at 1632 cm⁻¹ (C=O stretching of amide I), 1540 cm⁻¹ (N−H bending of amide II), and 1239 cm⁻¹ (C−N stretching coupled with N−H bending of amide III), confirming the successful conjugation of methacrylate groups and validating the effective methacrylation of gelatin.

The XRD patterns of gelatin and GelMA provide insight into the structural modifications induced by the methacrylation process (**Figure 2c**). Following methacrylation, the GelMA pattern exhibits a noticeable shift and disorders in peak intensity and position, suggesting alterations in molecular packing and crystallinity. The reduction in peak intensity in the GelMA spectrum implies a disruption in the hydrogen bonding network and structural rearrangement caused by the incorporation of methacrylate groups. This modification reduces the degree of molecular order, confirming the successful conjugation of methacrylate moieties onto the gelatin backbone.

Finally, FE-SEM analyses, encompassing both cross-sectional (**1**) and surface views (**2**), were employed to elucidate the structural characteristics of the synthesized GelMA (**Figure 2d**). The cross-sectional SEM image reveals a highly wrinkled and sheet-like morphology, indicative of the polymeric nature of GelMA. The presence of these thin, folded layers suggests that the material has retained a degree of flexibility, which may be influenced by the degree of methacrylation and the drying process. The interconnected sheet-like structures observed could play a role in subsequent hydrogel formation by facilitating efficient cross-linking and network formation when photopolymerized. In contrast, the surface morphology appears denser and more compact, with fewer visible folds or irregularities compared to the cross-sectional view. This difference may be attributed to polymer packing at the surface during solvent evaporation or the influence of processing conditions such as freeze-drying. The relatively smoother surface texture suggests a more homogeneous distribution of functional groups, which could impact the efficiency of photo-crosslinking and mechanical properties of the resulting hydrogel.

### 2.2. Effect of Photoinitiators on Stiffness of GelMA Hydrogels

The optimization of cross-linking conditions in GelMA hydrogels plays a crucial role in tailoring their physicochemical and biological properties by modulating their structural and mechanical characteristics. To date, while several studies, including our own, have explored the tuning of visible light-induced cross-linking parameters such as co-initiator or GelMA prepolymer concentration, these investigations have primarily focused on refining conditions for EY-based photopolymerization. ^23, 40, 41^ Despite their valuable contributions in guiding researchers toward optimal EY-mediated GelMA cross-linking conditions, a critical gap remains: the comparative assessment of different photoinitiator systems within the same biomaterial framework. To bridge this gap, we conducted a systematic and comprehensive study evaluating the impact of three distinct photoinitiators, EY, LAP, and Ru, at varying concentrations on the physicochemical and biological properties of 5% and 10% (w/v) GelMA hydrogels. Specifically, we examined their effects on **(i)** rheological properties, **(ii)** enzymatic degradation, **(iii)** swelling behavior, and **(iv)** cell viability, providing a systematic analysis on how different photoinitiation systems influence the crosslinked GelMA network.

#### 2.2.1. Effect of [EY] on Stiffness

The real-time rheological properties within the linear viscoelastic region (LVER), combined with the calculated Young’s modulus values, were examined to assess the degree of cross-linking, gelation profile, and the mechanical performance of the hydrogels. **Figure 3** and **Figure 4** summarize the impact of EY, LAP, and Ru concentrations in 5 and 10% (w/v) GelMA prepolymer solutions on the polymerized hydrogel’s stiffness, respectively.

**Figure 3.**
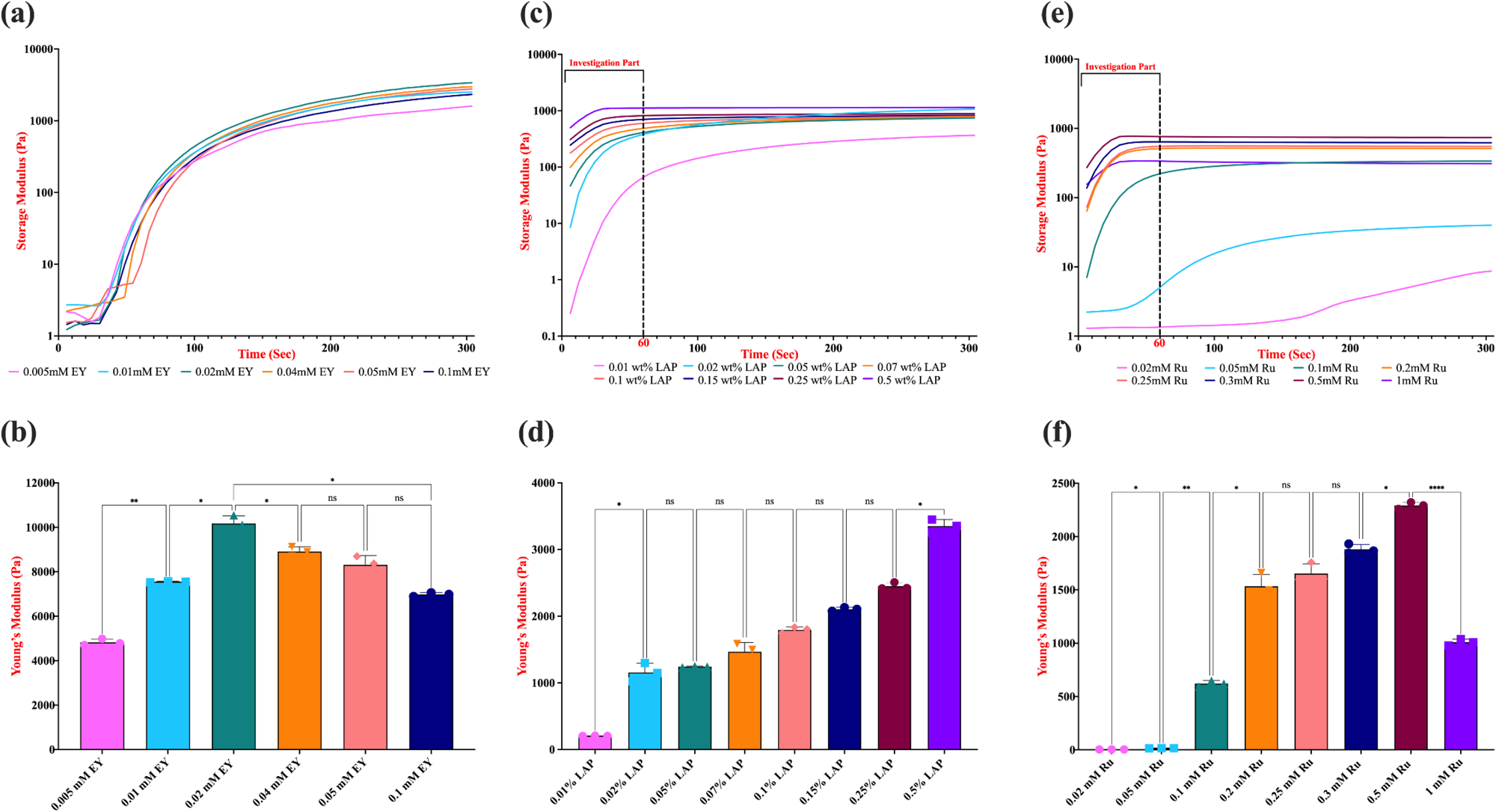
Effect of EY, LAP and Ru photoinitiator concentrations on stiffness. 5% (w/v) GelMA hydrogels were photocross-linked on a parallel plate rheometer at room temperature under specific light conditions (*λ=*530 nm for EY, 365 nm for LAP, and 430 nm for Ru), each at an intensity of 10 mW/cm^2^ for 5 minute to measure real-time Storage modulus and Young’s modulus of EY (**a, b**), LAP (**c, d**), and Ru (**e, f**) photoinitiating system. Prior to cross-linking, samples were incubated at 37°C. The data represent the average and smoothed storage modulus (*G′*) obtained from three independent replicates. Statistical analysis was conducted using a one-way ANOVA followed by Tukey’s post hoc test, with a significance threshold of p ≤ 0.05. Statistical notations are as follows: n.s. (not significant, p > 0.05), * (p ≤ 0.05), and **** (p ≤ 0.0001). Error bars shown indicate (n=3) the standard deviation.

**Figure 4.**
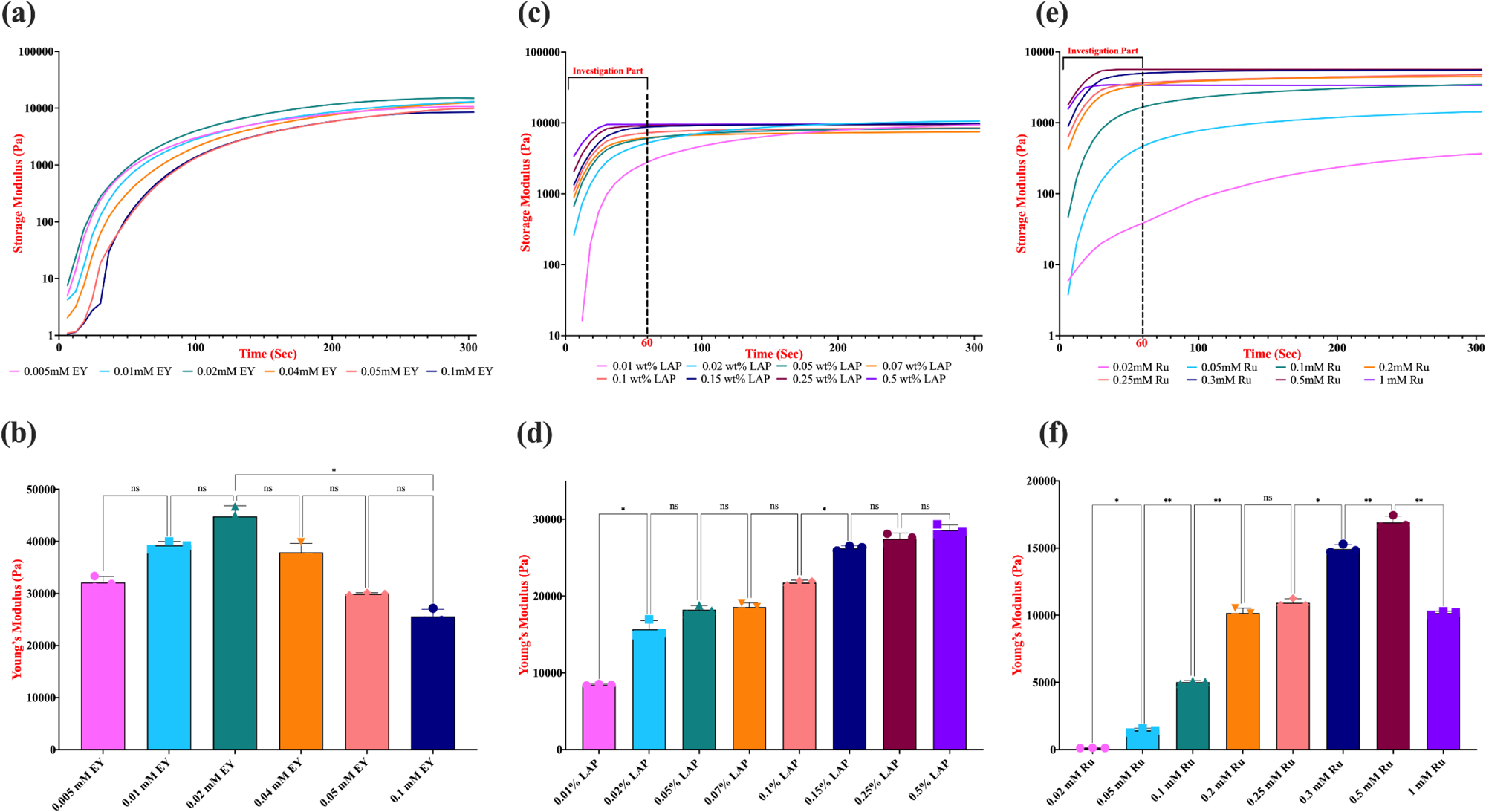
Effect of EY, LAP and Ru photoinitiator concentrations on stiffness. 10% (w/v) GelMA hydrogels were photo-crosslinked on a parallel plate rheometer at room temperature under specific light conditions (*λ=*530 nm for EY, 365 nm for LAP, and 430 nm for Ru), each at an intensity of 10 mW/cm^2^ for 5 minute to measure real-time Storage modulus and Young’s modulus of EY (**a, b**), LAP (**c, d**), and Ru (**e, f**) photoinitiating system. The data represent the average and smoothed storage modulus (*G′*) obtained from three independent replicates. Statistical analysis was conducted using a one-way ANOVA followed by Tukey’s post hoc test, with a significance threshold of p ≤ 0.05. Statistical notations are as follows: n.s. (not significant, p > 0.05), * (p ≤ 0.05), and **** (p ≤ 0.0001). Error bars shown indicate (n=3) the standard deviation.

In our previous study, we optimized the concentration of EY within the range of 0.005 to 0.5 mM in 5% (w/v) GelMA hydrogels and observed that concentrations up to 0.1 mM were highly cytocompatible on NIH-3T3 fibroblast cells. Nevertheless, at 0.2 mM and above, we observed a notable decline in cell viability, indicating cytotoxic effects. ^40^ Based on these findings, we decided to focus on investigating GelMA prepolymer solutions using [EY] from 0.005 to 0.1 mM (0.005, 0.01, 0.02, 0.04, 0.05, 0.1 mM), while maintaining constant co-initiator and comonomer concentrations (100 mM TEA and 50 mM NVP) and standardized light exposure parameters (10 mW/cm^2^, 5 minutes). It is important to note that this study does not aim to optimize the concentrations of TEA and NVP; rather, their levels were intentionally kept constant to isolate and evaluate the specific effect of photoinitiator concentration, [EY], on hydrogel stiffness and network properties.

Our studies demonstrated that the mechanical properties of the produced 5% GelMA hydrogels are strongly influenced by [EY] (**Figure 3a** and **3b**). Specifically, both the real-time storage and Young’s moduli values of hydrogels exhibited a direct correlation with EY concentrations below 0.02 mM, indicating that increasing [EY] within this range enhances the hydrogel’s stiffness from ∼4.8 kPa to ∼10.1 kPa. However, at EY concentrations exceeding 0.02 mM, this trend reverses, resulting in a decrease in stiffness below ∼7 kPa, which further confirms our previous findings. ^40^ Furthermore, real-time monitoring of the elastic modulus has enabled us to examine the gelation profile of these hydrogels, providing deeper insights into their cross-linking dynamics.

The mechanical behavior observed across different [EY] conditions can be directly attributed to the molecular mechanism of the EY/TEA/NVP photoinitiation system. Upon green light irradiation (λ ≈ 500–550 nm), EY transitions from its ground state (S₀) to an excited singlet state (S₁), which then undergoes intersystem crossing to a long-lived triplet state (T₁). In this excited triplet state, EY acts as a strong oxidant and accepts an electron from the tertiary amine TEA. This electron transfer results in the generation of an EY radical anion and a TEA-derived radical cation. The TEA radical cation undergoes deprotonation to yield α-amino alkyl radicals, which are the primary initiators that attack the carbon-carbon double bonds in methacryloyl groups, initiating free-radical polymerization. ^42^ In this system, NVP plays a dual role: **(i)** it acts as a reactive diluent, and **(ii)** a comonomer that readily copolymerizes with methacryloyl groups on GelMA, thereby improving network homogeneity and minimizing oxygen inhibition. ^40^ Additionally, NVP’s electron-rich vinyl group enhances the overall propagation rate by stabilizing the radical intermediates formed during chain growth.

At EY concentrations up to 0.02 mM, the generation of initiating radicals is efficient and balanced, enabling effective chain propagation and cross-linking. This is supported by the increase in both storage modulus and Young’s modulus within this range, indicating formation of a mechanically robust and homogeneously crosslinked hydrogel network. The gelation time, time required for the crossover point of G′ and G″, shortens due to effective radical initiation and propagation dynamics, reflecting rapid network formation.

However, beyond 0.02 mM EY, we observed a decline in stiffness, despite the continuation of fast gelation kinetics. This can be attributed to an overabundance of excited EY molecules, which may lead to non-productive photophysical processes, such as triplet-triplet annihilation and self-quenching. Additionally, excess radical species produced at high [EY] can accelerate bimolecular termination reactions, resulting in shorter polymer chains and incomplete cross-linking. These phenomena may lead to the formation of dangling chains, polymer strands that are covalently linked to the network at only one end, thereby decreasing effective crosslink density and mechanical strength. ^43, 44^ Our results showing lower modulus values where [EY] higher than 0.04 mM, despite rapid gelation, are consistent with this mechanistic understanding. Comparison of 5% and 10% (w/v) GelMA hydrogels under identical photoinitiating conditions further revealed a significant enhancement in stiffness with increasing polymer concentration (**Figure 3a, 3b** and **4a, 4b**). While the trends in mechanical performance and gelation kinetics relative to [EY] were consistent across both GelMA concentrations (G′ increases up to 0.02 mM [EY], G′′ decreases beyond 0.02 mM [EY]), the values of Young’s modulus were significantly higher in 10% GelMA hydrogels (∼10.1 kPa vs. ∼45 kPa for 0.02mM). This can be attributed to the increased availability of methacryloyl groups, which promotes more extensive cross-linking and results in a denser polymer network. Notably, 10% (w/v) GelMA hydrogels also exhibited faster gelation kinetics at equivalent [EY], suggesting that polymer concentration not only influences final mechanical properties but also impacts the dynamics of network formation.

In conclusion, our results demonstrate that both the concentration of photoinitiator and the GelMA polymer content are critical parameters in modulating hydrogel mechanics. The EY-photoinitiating system operates through a light-induced electron transfer mechanism that generates free radicals capable of initiating and propagating polymer chains. TEA serves as a radical donor, while NVP improves propagation kinetics and polymer network integrity. At optimal [EY], these components act synergistically to promote efficient, homogeneous cross-linking and produce stiff hydrogels. However, excessive [EY] disrupts this balance through photophysical quenching and radical overproduction, ultimately reducing mechanical performance. Moreover, increasing GelMA content enhances network density and stiffness, though at the cost of reduced porosity and molecular diffusivity. These findings highlight the importance of finely tuning photoinitiator chemistry and polymer concentration in designing hydrogels with desired mechanical and biological performance for specific biomedical applications.

#### 2.2.2. Effect of [LAP] on Stiffness

Next, in determining the range of [LAP] on properties of hydrogels, we referenced existing literature on the cytotoxicity of LAP-based photopolymerization systems, particularly in biomedical applications involving GelMA. Many studies have demonstrated that LAP concentrations exceeding 0.5% (w/v) exhibit cytotoxic effects across various cell types, including human renal proximal tubule endothelial cells, M-1 mouse collecting duct cells, 3T3 fibroblasts. ^21, 45 46^ Therefore, in our study, we focused on assessing the mechanical properties of GelMA hydrogels with [LAP] ranging from 0.01% to 0.5% (w/v) (specifically, 0.01, 0.02, 0.05, 0.07, 0.1, 0.15, 0.25, and 0.5% w/v). All experiments were conducted under standardized light exposure conditions (10 mW/cm^2^, 365 nm UV light, 5 min). Fairbanks *et al*. reported that LAP has a maximum molar extinction coefficient, 218 M⁻¹cm⁻¹, at 365 nm UV, and around 25 M⁻¹cm⁻¹ at 405 nm visible light. Their comparative analysis with 10 mW/cm^2^ intensity of light demonstrated that gelation of PEGDA under 365 nm light was six times more efficient than 405 nm at the same intensity, without cytotoxic effects. ^31^ Given this efficiency difference, we decided to utilize 365 nm UV light to investigate the effects of [LAP] on GelMA hydrogel properties more effectively. Our analysis confirmed that LAP system exhibits a rapid gelation profile, demonstrating a steep increase in storage modulus during the initial seconds of irradiation at 365 nm of light, especially exceeding 0.05% (w/v)% LAP concentration (**Figure 3c**). This trend indicates that LAP performs directly correlated photoinitiation efficiency and polymerization kinetics with increasing concentration, indicating a swift network formation between methacryoyl chains due to higher and faster free radical production.

To ensure consistency across all photoinitiating systems in terms of light exposure initiation, we applied uniform rheometer configurations to facilitate a direct comparison of their gelation profiles. In all three systems, the first real-time storage modulus measurement was recorded precisely at 12 seconds. This initial "lag time" included a 6-second light exposure period to allow for equilibrium in the rheometer analysis, followed by an additional 6 seconds before capturing the first data point. Our findings indicate that all GelMA prepolymer solutions, except the one containing 0.01% (w/v) LAP, experienced gelation within this 12-second period, highlighting the superior free radical production efficiency of the LAP system (**Figure 3c**). 10% (w/v) GelMA hydrogels formulated with the same LAP concentrations exhibited an even more rapid and efficient gelation profile, with all samples achieving gelation within 6 seconds. This accelerated cross-linking behavior can be attributed to the higher density of methacryloyl groups present in the more concentrated GelMA network, which enhances the availability of reactive sites for polymerization (**Figure 4c**).

Particularly, an interesting observation in our time-sweep oscillatory tests was the unexpected behavior of LAP-based photopolymerization in reaching its ultimate storage modulus. Many studies investigating UV-light-initiated photopolymerization, whether utilizing I2959 or LAP, have reported a direct correlation between increased photoinitiator concentration or UV light exposure and higher elastic or storage modulus of the hydrogel. ^47, 48, 49^ However, while our results are largely consistent with these findings, certain specific contradictions were observed. Even though it is anticipated that lower LAP concentrations, such as 0.02% (w/v), would yield a lower storage modulus, our data revealed an unexpected higher final storage modulus with 5% GelMA hydrogels at this concentration compared to all higher [LAP] values, except for 0.5% (w/v), under a 5-minute time-sweep oscillatory test. LAP concentrations higher than 0.05% (w/v) resulted in a stable viscoelastic response after approximately 1 minute of exposure to 365 nm light, indicating that the hydrogel network had reached its equilibrium state. In contrast, GelMA prepolymer solutions containing LAP concentrations below 0.05% (w/v) exhibited a continuous increase in storage modulus over time, suggesting ongoing network formation and progressive densification without achieving equilibrium (**Figure 3c**).

Typically, higher initiator concentrations generate a greater number of propagating sites, leading to shorter polymer chain lengths and an increased crosslink density per unit volume. As a result, the effective crosslink density is expected to increase (corresponding to a lower average molecular weight between crosslinks, *Mc*), ultimately leading to increased stiffness and reduced swelling ratio. In all free-radical polymerizations, the propagation rate of radicals follows first order kinetics with respect to their concentration, while their termination rate follows second order kinetics. ^50^ Consequently, increasing the initiator concentration accelerates radical termination. In UV-initiated polymerization, the overall reaction kinetics are known to favor an increased termination rate. ^51^ Based on this knowledge, we think that as the [LAP] rises, the increasing presence of free radicals through the high concentration (0.1% (w/v) or higher) may promote a greater frequency of premature chain termination events, resulting in an abundance of uncrosslinked methacryloyl chains. While some of these chains are completely washed out, others become partially incorporated into the network at only one end, forming dangling chains. These dangling chains may contribute to an increased excluded volume within the GelMA network, potentially explaining the deviation from conventional photopolymerization polymerization behavior. ^52^ To summarize, the observation that 0.02% (w/v) LAP ultimately yields a higher storage modulus than higher concentrations of LAP suggests that a slower and more efficient cross-linking process, characterized by fewer radical termination events and greater network homogeneity, results in a mechanically superior hydrogel. In contrast, at higher LAP concentrations (e.g., 0.5% (w/v)), excessive radical formation significantly accelerates gelation but compromises the integrity of the final crosslinked structure.

We further confirmed this atypicality of LAP photoinitiating system with 10% (w/v) GelMA prepolymer solutions under identical conditions. This time, we more clearly observed higher ultimate storage modulus with both lower concentrations, 0.01 and 0.02% (w/v), than all the superior [LAP], indicating the interesting photopolymerization kinetics of varying concentrations of this photoinitiating system to produce GelMA hydrogels (**Figure 4c**). Similar discrepancies have been reported by Nguyen *et al.* and further substantiated by Connell *et al.*, who concluded that UV light intensity and photoinitiator concentration do not solely determine the final storage modulus but instead influence the cross-linking rate and efficiency. ^21, 12^

To systematically analyze the influence of varying photoinitiator concentrations on the mechanical properties of GelMA hydrogels, it was essential to ensure that the hydrogels achieved stable gelation while maintaining comparable stiffness characteristics. Given the rapid gelation profiles of photoinitiators such as LAP and Ru, we selected a 1-minute exposure period for our further investigations. This approach minimized the atypical behaviors observed at low photoinitiator concentrations, which can otherwise lead to misinterpretations of the hydrogels’ Young’s modulus, typically derived from the final measured storage modulus data. Following this approach, the Young’s moduli values of 5% (w/v) GelMA hydrogels crosslinked with varying LAP concentrations demonstrated a clear positive correlation between [LAP] and hydrogel stiffness (**Figure 3d**). The highest Young’s modulus was observed at approximately ∼3.3 kPa for hydrogels polymerized with 0.5% (w/v) LAP, indicating an enhanced cross-linking density and mechanical strength of the network. A comparable trend was observed in 10% (w/v) GelMA hydrogels crosslinked with the same range of LAP concentrations, with the highest stiffness of ∼29 kPa (**Figure 4d**). Notably, due to the higher availability of methacrylate groups for cross-linking, the effectiveness of lower LAP concentrations, such as 0.15% and 0.25% (w/v), appeared to be enhanced. This was reflected in the absence of a statistically significant difference in Young’s moduli values between these conditions and the highest [LAP], suggesting a saturation effect in cross-linking efficiency at elevated GelMA concentrations.

When evaluating the Young’s moduli of 5% and 10% (w/v) GelMA hydrogels crosslinked with varying [LAP] concentrations after 5-minutes of exposure to 365 nm UV light rather than 1-minute, the previously noted atypical behavior of LAP became more pronounced. Specifically, the trend of achieving a higher Young’s modulus at lower LAP concentrations such as 0.01 and 0.02% (w/v) compared to higher concentrations was more evident (∼32 kPa vs ∼29 kPa), highlighting the influence of LAP’s unique photopolymerization mechanism under UV light conditions owing to changing amount and rate of free radical production. (**Figure S1**) Although our study did not extend to GelMA concentrations exceeding 10% (w/v), we anticipate that the asymptotic behavior observed in LAP-mediated photopolymerization with UV light would become even more pronounced at higher concentrations. Further investigations into LAP’s unique free radical photopolymerization mechanism under varying polymerization conditions are required to fully understand and optimize hydrogel performance for biomedical applications.

#### 2.2.3. Effect of [Ru] on Stiffness

We further extended our analysis of the photopolymerization system for GelMA hydrogels by exploring the Ru/SPS photoinitiating system, which has gained significant attention in the field. Our investigation range for this system was determined based on an extensive review of the literature, which commonly reports the use of Ru concentrations between 0.1 and 2 mM, and a fixed 1:10 molar ratio (Ru:SPS) with different type of biomaterials. Although several studies, which demonstrates the applicability and advantageous characteristics of Ru-mediated photopolymerization especially on 3D bioprinting systems, were published over the years ^25, 26, 35, 53^, to the best of our knowledge, there are no specific studies for investigating the [Ru] on mechanical properties of GelMA hydrogels since the first application of Ru/SPS visible-light photoinitating system for cross-linking of gelatin-based materials. ^54^ Thus, with this study, we decided to explore the effect of Ru concentration on 5% and 10% (w/v) GelMA hydrogels across a range of 0.02 to 1 mM (specifically, 0.02, 0.05, 0.1, 0.2, 0.25, 0.3, 0.5, and 1 mM), while maintaining a fixed [SPS] at a 1:10 ratio (Ru:SPS). Our real-time rheological analysis revealed that increasing [Ru] enhances the photopolymerization rate and cross-linking efficiency, with a notable increase of storage modulus up to 0.5 mM (**Figure 3e**). In contrast, lower concentrations, including 0.02 mM and 0.05 mM, failed to induce substantial gel formation in 5% (w/v) GelMA content. Similar to the LAP photoinitiating system, gelation in 5% and 10% (w/v) GelMA prepolymer solutions with Ru/SPS system was observed within the first 5 to 10 seconds of 430 nm blue light irradiation (**Figure 3e** and **4e**), likely due to the exceptionally high molar extinction coefficient of Ru (12391 M.cm^-1^ at 430 nm). ^55^

To enable a more accurate comparative analysis of the LAP and Ru photoinitiating systems, we employed the previously described approach to evaluate the overall stiffness profiles under varying conditions for the Ru-based system. Specifically, we focused on photorheological analysis within the first minute of irradiation, using the ultimate storage modulus at the 1-minute mark to construct Young’s modulus graphs. Corresponding stiffness profiles clearly demonstrated the exact positive correlation of varying concentrations of Ru in both 5% and 10% GelMA hydrogels up to 0.5 mM level (**Figure 3f** and **4f**). However, at Ru concentrations exceeding 0.5 mM, such as 1 mM, the hydrogels exhibited diminished mechanical integrity causing a reduction in stiffness of resulting hydrogels. Also, at concentrations beyond 1 mM, pre-gelation occurred even before exposure to 430 nm blue light (data not shown). This phenomenon can be attributed to the high reactivity of the Ru/SPS system with high concentrations (1:10 or higher Ru:SPS molar ratio) in visible light range, leading to premature cross-linking before rheological measurements started and compromise hydrogel properties.

An intriguing observation was the more pronounced transition in Young’s modulus from 1-minute to 5-minute light exposure in 10% (w/v) GelMA hydrogels, particularly within the 0.2 to 0.5 mM Ru range (**Figure S2**). While no significant differences were detected in the Young’s modulus of 5% (w/v) GelMA hydrogels between 1-minute and 5-minute exposure to 430 nm light, the 10% (w/v) GelMA hydrogels exhibited a distinct trend. With extended light exposure, the stiffness differences between lower Ru concentrations (<0.5 mM) progressively diminished, as these conditions continued to form a denser cross-linked network. In contrast, at 0.5 mM Ru, cross-linking appeared to reach saturation, resulting in a final stiffness of approximately 17 kPa, aligning with that of lower concentrations after prolonged irradiation. Overall, this observed convergence in Young’s modulus values for 10% (w/v) GelMA hydrogels under varying Ru concentrations and exposure times suggests that, beyond a certain initiator concentration and irradiation period, the network reaches a saturation point where additional cross-linking does not significantly enhance stiffness. Comparable patterns at elevated concentrations have also been reported and discussed for the LAP-based photoinitiating system in previous section. This plateau effect indicates that optimal mechanical properties can be achieved without prolonged exposure or excessive initiator use to prevent potential cytotoxicity, which is advantageous for biomedical applications.

The Ru/SPS co-initiator system uniquely operates through a photoredox mechanism, whereas LAP and EY systems primarily rely on direct radical generation upon light activation, without involving a redox cycle with a co-initiator. Upon exposure to visible light, Ru^2+^ becomes photoexcited and oxidizes to Ru^3+^ by transferring electrons to SPS. This electron transfer causes SPS to dissociate into sulfate anions and highly reactive sulfate radicals. These sulfate radicals have been shown to facilitate step-growth thiol-ene polymerization in gelatin-based hydrogels. ^53, 56^ In GelMA, dissociated sulfate radicals initiate chain-growth polymerization of MA groups in seconds, forming nondegradable oligomethacryloyl kinetic chains that crosslink the hydrogel network. ^35^ In 2021, Yang *et al*. highlighted that modifying SPS concentrations is an effective strategy to tailor the mechanical properties of GelAGE hydrogels, owing to the unique photo-redox mechanism of the Ru/SPS photoinitiating system. ^56^ However, there is lack of studies in literature presenting the impact of [SPS] on free radical photopolymerization systems, such as GelMA. Therefore, in our current research, we also explored how varying [Ru]:[SPS] molar ratios (specifically 1:1, 1:2, 1:5, 1:10 (our standard condition), 1:15, and 1:20) affect the viscoelastic properties and stiffness of 5% and 10% (w/v) GelMA hydrogels. For this investigation, we maintained constant Ru concentrations of 0.2 mM and 0.3 mM for the 5% and 10% (w/v) GelMA prepolymer solutions, respectively. Our results revealed that adjusting the [SPS] molar ratio is pivotal in the Ru/SPS photoinitiating system for effective cross-linking of MA groups in GelMA hydrogels (**Figure S3**). Notably, increasing the Ru:SPS molar ratio up to 1:15 enhances hydrogel stiffness from approximately 1.5 to 2.2 kPa in 5% (w/v) GelMA hydrogels, and from about 13.5 to 24.5 kPa in 10% (w/v) GelMA hydrogels. However, further increasing the ratio to 1:20 did not yield significant improvements, suggesting a saturation point in sulfate radical production for effective cross-linking. Additionally, we observed that a minimum Ru:SPS molar ratio of 1:5 is necessary to achieve proper gelation of GelMA prepolymer solutions, as the chain-growth polymerization process is directly dependent on the availability of sulfate radicals. We believe that these identifications are particularly advantageous for biomedical applications with Ru/SPS photoinitiated systems, as it enables the tailoring of hydrogel mechanical properties without introducing excessive [SPS], which could potentially lead to cytotoxicity.

### 2.3. Effect of Photoinitiators on Enzymatic Biodegradation Profile of GelMA Hydrogels

To further confirm the impact of varying concentrations of EY, LAP and Ru photoinitiators on hydrogel stiffness, we conducted *in vitro* enzymatic biodegradation experiments using Collagenase from *Clostridium histolyticum*. Hydrogels were incubated in a 0.1% collagenase solution at 37 °C, with gravimetric measurements taken periodically over a 360-minute period (0, 15, 30, 45, 60, 75, 90, 135, 195, 255, 360 min) to monitor degradation until complete hydrogel dissolution. It is known from literature that GelMA hydrogels with higher stiffness typically exhibit slower enzymatic degradation due to their densely crosslinked networks, which hinder enzyme infiltration and substrate accessibility. This limited penetration is known to restrict the enzyme’s ability to interact with and degrade the GelMA’s enzyme-sensitive chains, thereby prolonging the hydrogel’s structural integrity. ^57^

Our gel retention findings for each of photoinitiating system align with discussed rheological analyses, indicating that the hydrogels with the highest stiffness exhibited the slowest degradation due to their densely crosslinked network, which restricts collagenase enzyme diffusion and penetration throughout the matrix (**Figure 5**). To be more specific, we observed that 5% (w/v) GelMA hydrogels crosslinked with 0.1 mM EY degraded completely within 135 minutes, a faster degradation rate compared to hydrogels prepared with 0.01- or 0.02-mM EY (**Figure 5a**). This finding supports our current rheological analyses and corroborates previous studies, which have demonstrated that increasing [EY] beyond optimal levels can lead to decreased mechanical properties and accelerated degradation due to over-initiated polymerization resulting in a less uniform network structure. ^40^ A similar degradation trend was observed in 10% (w/v) GelMA hydrogels, which exhibited longer degradation times compared to their 5% counterparts due to increased stiffness, denser cross-linking and reduced pore sizes, thereby enhanced resistance to enzymatic degradation (**Figure 5b**). ^58^

**Figure 5.**
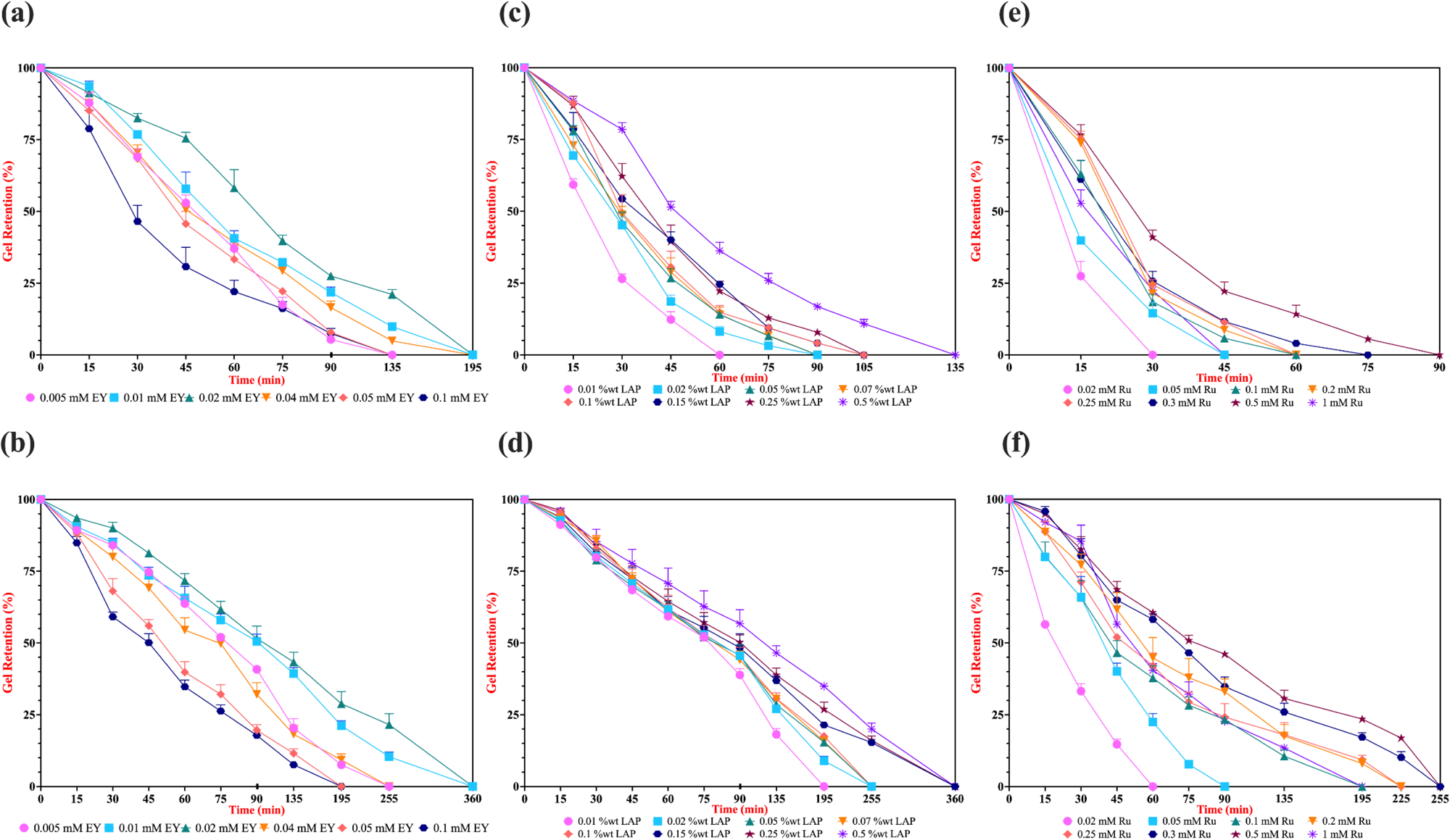
Effect of EY, LAP and Ru photoinitiator concentrations on enzymatic biodegradation profile. Hydrogels were photo-crosslinked in a 96-well plate at room temperature using hand-made light sources emitting at 530 nm for EY (5-minute exposure), 365 nm for LAP, and 430 nm for Ru (both with 1-minute exposures), each at an intensity of 10 mW/cm^2^. *In vitro* enzymatic degradation was assessed by incubating the hydrogels in a 0.1% collagenase solution until complete degradation. Panels (**a**) and (**b**) display the degradation profiles for EY-initiated systems, (**c**) and (**d**) for LAP-initiated systems, and (**e**) and (**f**) for Ru-initiated systems, corresponding to 5% and 10% (w/v) GelMA hydrogels, respectively. Error bars represent the standard deviation (n=3).

As previously-mentioned, the investigation for varying concentrations of LAP- and Ru-mediated photo-crosslinking on mechanical properties of GelMA hydrogels were standardized with 1-minute exposure of light for all corresponding prepolymer solutions to facilitate a more accurate comparison of these rapidly-acting photoinitiators. In the case of LAP-crosslinked 5% GelMA hydrogels (**Figure 5c**), degradation progressed rapidly at lower LAP concentrations, particularly at 0.01 and 0.02% (w/v), where complete degradation occurs within approximately 60 to 90 minutes. Hydrogels cross-linked with 0.05–0.25% (w/v) LAP showed slightly improved stability, degrading entirely between 90 and 120 minutes. At 0.5% (w/v) LAP, degradation was noticeably slower, with hydrogels retaining structural integrity for up to 135 minutes before complete breakdown. A similar pattern was observed in 10% (w/v) GelMA (**Figure 5d**), where overall degradation occurred at a significantly slower rate due to increased extent of polymer network. Hydrogels crosslinked with LAP concentrations up to 0.15% (w/v) withstood collagenase degradation for approximately 250 minutes, while higher concentrations (0.15, 0.25, and 0.5% (w/v)) prolonged stability up to 360 minutes. This trend completely aligns with our Young’s modulus analysis, including statistical significance assessments, further reinforcing that increased cross-linking density enhances resistance to enzymatic degradation, thereby improving hydrogel durability.

In Ru-crosslinked 5% GelMA hydrogels (**Figure 5e**), degradation was observed much faster, particularly at lower Ru concentrations. Hydrogels cross-linked with 0.02 and 0.05 mM [Ru] degrade entirely within 30 and 45 minutes, highlighting the insufficient network formation or larger pore sizes at these concentrations. Increasing [Ru] to 0.3 mM or even 0.5 mM improved structural integrity, extending hydrogel stability up to 75 and 90 minutes before complete dissolution, respectively. However, degradation rates of hydrogels cross-linked with 1 mM Ru were unexpectedly accelerated, with complete breakdown occurring within 45 minutes, like lower Ru conditions. This anomaly aligns with swelling and rheological results, where excessive Ru concentrations disrupt homogeneous network formation, leading to increased water or enzyme uptake and structural weaknesses. The effect is more pronounced in 10% GelMA (**Figure 5f**), where hydrogels at lower Ru concentrations (0.02 and 0.05 mM) degraded within 90 minutes, while intermediate concentrations (0.2–0.5 mM) extended stability to approximately 225 minutes. Even though condition with 1 mM [Ru] has shown a more stable profile compared to 5% (w/v) GelMA hydrogels, degradation accelerated again, with complete hydrogel loss occurring before 195 minutes, supporting our hypothesis that excessive Ru leads to network defects that make the hydrogel more prone to enzymatic breakdown.

Comparing the enzymatic degradation or swelling profiles of GelMA hydrogels cross-linked using different photoinitiators is inherently complex due to variations in the resulting network architectures and photopolymerization efficiency of the system. However, as our study explores the impact of different photoinitiating systems (EY, LAP, and Ru) on the mechanical properties of a consistent GelMA hydrogel network, not only in a concentration-dependent manner but also through comparative analysis across these systems, we could draw conclusions based on our experimental findings. It is well-established in the literature that hydrogels, whether composed of the same or different materials, can exhibit diverse degradation behaviors. These behaviors may involve degradation through hydrolysis of the polymer backbone or through cleavage of the crosslinker itself, leading to varying resistance to dissolution. ^59^ Nevertheless, since GelMA inherently degrades through its matrix metalloproteinase (MMP) sites under the action of MMP-1, MMP-8, or MMP-13 collagenases, these enzymes are commonly utilized in degradation studies, highlighting hydrolysis of the polymer backbone as the primary degradation mechanism. ^60^ With this assumption of a consistent degradation pathway, we can qualitatively assess the structural differences among GelMA hydrogels cross-linked using various photoinitiator systems. Notably, despite exhibiting lower stiffness, hydrogels cross-linked with the LAP photoinitiation system demonstrated degradation profiles comparable to or longer than those cross-linked with the EY system (**Figure 5b vs. 5d**). Conversely, hydrogels cross-linked with the Ru system degraded most rapidly (**Figure 5f**), likely due to the distinct photoredox reaction mechanism of the Ru/SPS system causing lower overall stiffness characteristics on GelMA hydrogels. These observations confirm that beyond initial stiffness, factors such as network architecture and the type of cross-linking density (homogeneous or heterogeneous) play critical roles in determining the enzymatic degradation rates of these hydrogels.

To gain insight into the network architecture of 5% (w/v) GelMA hydrogels cross-linked using EY, LAP, and Ru-mediated systems under comparable stiffness conditions (0.01 mM EY, 0.15% (w/v) LAP, and 0.3 mM Ru), we examined their intrinsic microstructure through FE-SEM imaging at 200× and 500× magnifications (**Figure S4**). The images reveal notable variations in pore size, shape, and connectivity, which directly influence the mechanical and transport properties of the hydrogels. Hydrogels cross-linked with EY (**Figure S4a** and **S4b**) exhibited a highly porous and interconnected network, characterized by relatively uniform and small pores. This suggests that EY-based cross-linking facilitated homogeneous polymerization, leading to a well-structured network with balanced mechanical integrity and permeability. In contrast, LAP-crosslinked hydrogels (**Figure S4c** and **S4d**) displayed larger, more irregularly shaped pores. This observation aligns with the known rapid polymerization kinetics of LAP, which may cause localized cross-linking heterogeneity, resulting in variable pore structures. The increased pore size may enhance nutrient diffusion and swelling capacity but could compromise mechanical stability. The FE-SEM images of Ru-crosslinked GelMA hydrogels (**Figure S4e** and **S4f**) indicated that the pore walls appear thinner and more irregular, with some regions showing collapsed or partially degraded structures. This confirms that the hydrogel network formed via Ru-mediated cross-linking have lower crosslink density or weaker polymer connectivity, leading to reduced mechanical robustness.

Therefore, it can be deduced that when selecting a photoinitiator system for GelMA hydrogel fabrication, it is crucial to consider not only the mechanical properties but also the degradation characteristics, especially for applications in tissue engineering and regenerative medicine.

### 2.4. Effect of Photoinitiator on Swelling Profile of GelMA Hydrogels

The swelling properties of hydrogels are critical for biomedical applications as they influence nutrient diffusion, waste removal, and overall cellular activities within the scaffold. The ability of a hydrogel to retain water is inversely correlated with its cross-linking density, where highly crosslinked networks exhibit lower swelling due to reduced free volume. ^61^ Therefore, to further assess the mechanical properties of resulting GelMA hydrogels cross-linked with different type and concentration of photoinitiators, we have conducted swelling studies on each hydrogel. Our findings on the swelling behavior of EY, LAP and Ru photoinitiating systems across varying concentrations further reinforce our rheological analysis. Specifically, conditions exhibiting higher Young’s modulus displayed lower swelling, which can be attributed to a more densely crosslinked network and reduced pore size. (**Figure 6**)

**Figure 6.**
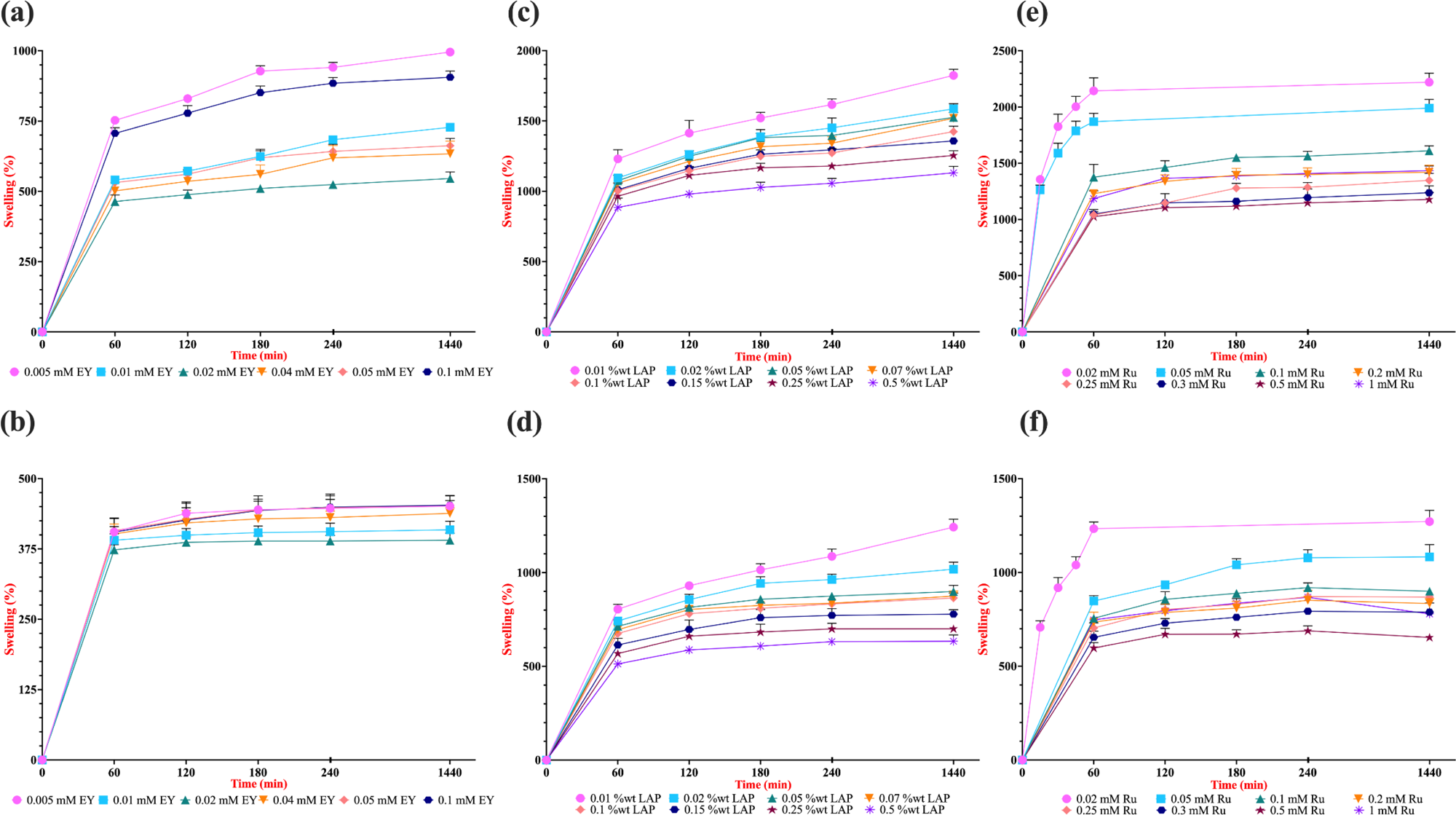
Effect of EY, LAP and Ru photoinitiator concentrations on swelling profile. Hydrogels were photo-crosslinked in a 96-well plate at room temperature using custom-built light sources emitting at 530 nm for EY (5-minute exposure), 365 nm for LAP, and 430 nm for Ru (both with 1-minute exposures), each at an intensity of 10 mW/cm^2^. Swelling percentage was assessed by incubating the lyophilized hydrogels in a PBS solution. Panels (**a**) and (**b**) display the swelling profiles for EY-initiated systems, (**c**) and (**d**) for LAP-initiated systems, and (**e**) and (**f**) for Ru-initiated systems, corresponding to 5% and 10% (w/v) GelMA hydrogels, respectively. Error bars represent the standard deviation (n=3).

In the EY-mediated system, 5% (w/v) GelMA hydrogels crosslinked with 0.02 mM EY exhibited approximately 550% swelling after one day of incubation in PBS, while hydrogels containing 0.04-, 0.05-, and 0.1-mM EY demonstrated swelling ratios of 633%, 662%, and 905%, respectively (**Figure 6a**). Once again, these results support our findings that beyond a certain [EY], the mechanical integrity of the hydrogel begins to deteriorate. Although the differences among these conditions were relatively small due to the highly crosslinked network and increased stiffness, a similar trend was also observed in 10% (w/v) GelMA hydrogels (**Figure 6b**).

Next, 5% (w/v) GelMA hydrogels cross-linked using varying concentrations of LAP exhibited swelling ratios ranging from approximately 900% to 1200% after just one hour of incubation in PBS, indicative of either highly porous, or heterogeneous network architecture (**Figure 6c**). Obtained findings corroborate our rheological analysis, which demonstrated a concentration-dependent stiffness profile within the initial 60 seconds of 365 nm UV light irradiation, with the lowest swelling (∼1130% at 24 hours) observed at 0.5% (w/v) LAP concentration. Our continued analysis with 10% (w/v) GelMA hydrogels cross-linked with decreasing LAP concentrations showed similar but less increase of swelling profile (**Figure 6d**). After 1 minute of light irradiation, hydrogels prepared with the lowest [LAP] demonstrated a swelling ratio of approximately 1240% following 24 hours of incubation in PBS, conversely, those synthesized with the highest [LAP] swelled to about 630% of their dry weight under identical conditions. These observations are consistent with the understanding that higher concentrations of GelMA naturally promote increased cross-linking between methacryloyl groups, resulting in a denser network structure and diminished swelling capacity. ^62, 63^ Additionally, our observed swelling ratios for both 5% and 10% (w/v) GelMA hydrogels photopolymerized using the LAP system align closely with findings from a recent study that examined the mechanical properties of 5%, 10%, and 15% (w/v) GelMA hydrogels crosslinked with 0.1% (w/v) LAP under 5 mW/cm^2^, 365 nm UV light for 3 minutes, a protocol quite similar to ours. ^64^ This concordance further validates the accuracy of our results within this system despite our decisional 1-minute of cross-linking and investigation of mechanical properties.

However, we hypothesize for our investigation that extending the duration of cross-linking of hydrogels with UV light irradiation to 5 minutes or longer, would reveal swelling and enzymatic degradation profiles less directly dependent on increasing [LAP]. This effect is likely due to the unique free radical photopolymerization mechanism of LAP under prolonged UV light exposure, as we discussed in **Figure S1**. Specifically, the hydrogels synthesized with 0.01, 0.02, or 0.05% (w/v) LAP would exhibit the least swelling under extended photopolymerization conditions, originating from a stiffer, less dangled network. We think that this phenomenon is not solely attributed to use of LAP but is influenced by the behavior of MA groups attached to gelatin chains during UV irradiation. Notably, we observed that extended UV exposure (beyond 3 minutes) might have the tendency to invert typical swelling and degradation characteristics, leading to increased swelling with higher LAP concentrations (data not shown). Similar atypical swelling behaviors have been observed before with copolymeric hydrogels of NVP and 2-hydroxyethyl methacrylate (HEMA), where oxidative decomposition during UV irradiation—due to increased UV intensity or exposure time—was discussed as responsible for more dangling chains within the network. ^65^ Furthermore, higher photoinitiator concentrations can accelerate gelation rates as discussed before, increasing chain termination events and resulting in more dangling chain ends. These unincorporated or partially integrated chains (potential structural anomalies illustrated in **Figure S6**) may contribute to altered swelling and degradation profiles, ultimately accounting for the atypical behavior observed in GelMA hydrogels cross-linked with the LAP system under prolonged UV exposure.

Finally, the swelling behavior of Ru/SPS-crosslinked GelMA hydrogels reveals a clear concentration-dependent trend influenced by both GelMA content and [Ru]. In 5% (w/v) GelMA hydrogels (**Figure 6e**), lower Ru concentrations (0.02–0.05 mM) lead to significantly higher swelling, with a peak swelling of approximately 2500% observed at 0.02 mM Ru after 24 hours. This confirms that at low Ru concentrations, the cross-linking density is insufficient to form a tightly packed hydrogel network, allowing for greater water uptake. As the [Ru] increases, the swelling ratio gradually decreases, reaching ∼1100% at 0.5 mM Ru, indicative of a more densely interchain structure with reduced pore size compared to other conditions. Notably, the anomalously high swelling at 1 mM [Ru] can be attributed to excessively fast free radical production, leading to premature termination, inhomogeneous network formation, and macro void development, as we discussed in our rheological analyses. On the other hand, the sharp increase in swelling within the first 60 minutes indicates rapid hydration and network expansion, especially at lower Ru concentrations. This necessitated measuring swelling percentages within the first hour; beyond this timeframe, the hydrogels were unable to maintain structural integrity in PBS due to excessive water uptake leading to dissolution. This phenomenon, known as degradation-induced swelling, occurs when the polymer network loses its elastic restorative force, permitting further volume increase even after reaching equilibrium swelling. ^61^ A similar trend is observed in 10% (w/v) GelMA hydrogels (**Figure 6f**), though the extent of swelling is notably lower compared to the 5% GelMA samples. The highest swelling (∼1300%) occurred at 0.02 mM Ru, while the lowest swelling (∼650%) was seen at 0.5 mM Ru after 24 hours of incubation. This reduction in overall swelling compared to the 5% GelMA hydrogels can be attributed to, once again, the inherently higher cross-linking density in the more concentrated GelMA network, which restricts water penetration and expansion. ^63^

As we discussed before, comparing the swelling characteristics of 5% and 10% (w/v) GelMA hydrogels cross-linked with EY, LAP, and Ru photoinitiating systems presents complexities due to inherent variations in the resulting network architectures. Nevertheless, such an analysis is essential to gain insights into the structural nuances of these networks as variations in swelling behavior directly influence the physical properties of the hydrogels, ultimately determining their suitability for different biomedical applications. ^66^ Our swelling analyses indicate that cross-linking density, inferred from the stiffness of the hydrogels (**Figure 3** and **Figure 4**), plays a pivotal role in determining their swelling capacity. For instance, 5% (w/v) GelMA hydrogels cross-linked with the EY-photoinitiating system exhibited the highest stiffness, leading to the lowest swelling profile (∼550% at its minimum), whereas those polymerized with LAP and Ru systems demonstrated significantly higher swelling capacities, with minimum values of approximately ∼1130% and ∼1200%, respectively (**Figure 6**). While Young’s modulus is a key parameter for interpreting swelling and enzymatic degradation profiles, stiffness alone does not fully explain the behavior of hydrogels cross-linked with LAP and Ru systems. Notably, in 10% (w/v) GelMA hydrogels, despite LAP-based networks exhibiting superior resistance to enzymatic degradation compared to Ru-based hydrogels, an intriguing trend emerged. The lowest swelling conditions for both Ru- and LAP-crosslinked hydrogels showed nearly identical swelling percentages (∼650%), despite a significant stiffness difference (∼30 kPa vs. 17 kPa). This suggests that the LAP-photopolymerized network possesses a unique water retention capability, likely influenced by its interesting methacryloyl chain architecture through UV light irradiation.

The investigation of network architecture of different 5% (w/v) GelMA hydrogels cross-linked with EY-, LAP- and Ru- photoinitiating systems with FE-SEM images could be also an explanatory way for this investigation of swelling profile between Ru and LAP systems (**Figure S4**). Specifically, LAP-crosslinked hydrogels (**Figure S4c,** and **S4d**) exhibited larger and more irregular pores, a characteristic attributed to the rapid polymerization kinetics of LAP, which can lead to heterogeneous cross-linking. These larger pores probably increase the hydrogel’s capacity to retain water, aligning with the unexpectedly high swelling ratio observed despite its higher stiffness compared to Ru-crosslinked networks. In contrast, Ru-crosslinked hydrogels (**Figure S4e,** and **S4f**) showed thinner and more irregular pore walls, with regions appearing collapsed or partially degraded. This fragmented structure suggests a lower crosslink density and weaker polymer connectivity, leading to a less stable network that facilitates greater water uptake and expansion. The combination of these morphological differences explains why Ru-mediated hydrogels exhibit the highest swelling, while LAP-crosslinked hydrogels, despite their higher stiffness, still demonstrate significant swelling due to their unique pore architecture and water retention properties.

Swelling behavior is a critical parameter in the biomedical applications of GelMA hydrogels, as it directly influences their ability to retain bioactive molecules, facilitate cell migration, and regulate nutrient diffusion in tissue engineering, wound healing, and drug delivery systems. In our study, the swelling profiles of EY-, LAP-, and Ru-crosslinked GelMA hydrogels exhibit distinct characteristics that impact their functional applications. Ru-crosslinked hydrogels demonstrated the highest swelling due to their weaker polymer network and thinner pore walls, which could be advantageous for applications requiring rapid hydration and enhanced drug release but may compromise mechanical stability. LAP-crosslinked hydrogels, despite their higher stiffness, exhibited unexpectedly high swelling, likely due to their large and irregular pores, making them suitable for applications balancing mechanical integrity with moderate swelling. EY-crosslinked hydrogels, with their highly porous and interconnected structure, seemed to have a more controlled swelling behavior, which is beneficial for tissue engineering applications requiring a balance between hydration and stability. Tailoring the swelling properties of these hydrogels by selecting appropriate photoinitiators and appropriate concentrations allows for the development of optimized biomaterials suited for specific biomedical applications, where controlled expansion, structural integrity, and hydration are key factors in performance. ^66^

### 2.5. Effect of Photoinitiator on Cell Viability of GelMA Hydrogels

To evaluate the cytocompatibility of photopolymerized GelMA hydrogels, NIH-3T3 fibroblasts were seeded onto hydrogels crosslinked with varying concentrations of EY-, LAP-, and Ru-photoinitating systems. Cell survival was quantified after 7 days of culture and is presented in **Figure 7** for both 5% (**Figure 7a, c, e**) and 10% (**Figure 7b, d, f**) GelMA formulations.

**Figure 7.**
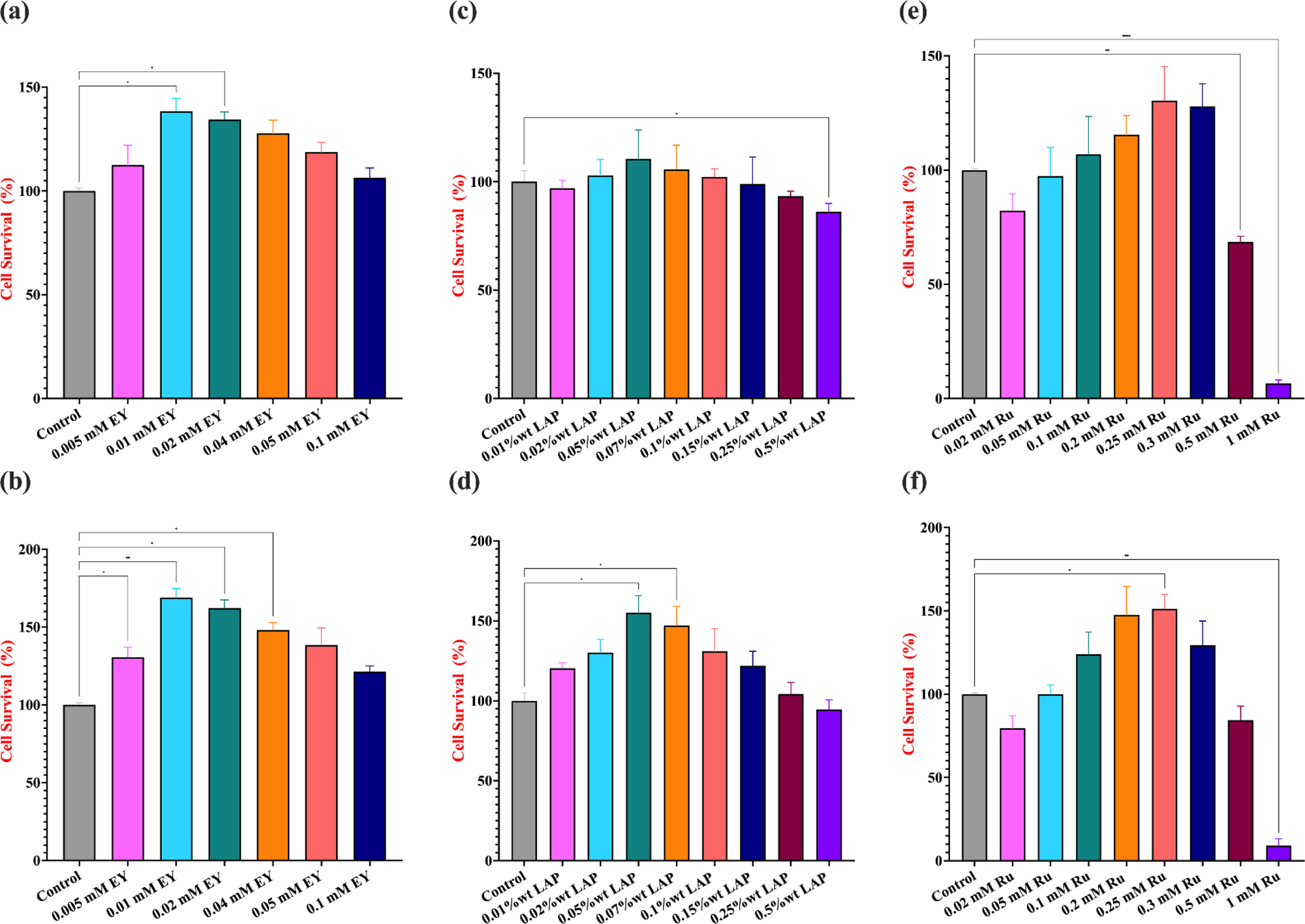
Effect of EY, LAP and Ru photoinitiator concentrations on cell viability. Hydrogels were photo-crosslinked in a 96-well plate at room temperature using custom-built light sources emitting at 530 nm for EY (5-minute exposure), 365 nm for LAP, and 430 nm for Ru (both with 1-minute exposures), each at an intensity of 10 mW/cm^2^. Cell viability was assessed by seeding NIH-3T3 cells on top of the GelMA hydrogels and cultured for 7 days. Graphs (**a**) and (**b**) display the cell survival for EY-initiated systems, (**c**) and (**d**) for LAP-initiated systems, and (**e**) and (**f**) for Ru-initiated systems, corresponding to 5% and 10% (w/v) GelMA hydrogels, respectively. Statistical analysis was conducted using a one-way ANOVA followed by Tukey’s post hoc test, with a significance threshold of p ≤ 0.05. Statistical notations are as follows: n.s. (not significant, p > 0.05), * (p ≤ 0.05), and **** (p ≤ 0.0001). Non-significant analyses were not demonstrated. Error bars represent the standard deviation (n=3).

EY is a non-toxic initiator, which can be excited with visible light (530 nm), making EY suitable for biological applications. ^67, 68^ In this study, the highest cell survival was observed at 0.01 mM EY, with a significant increase compared to lower (0.005 mM) and higher (≥0.04 mM) concentrations (**Figures 7a and 7b**). A dose-dependent decline in cell viability was apparent as EY concentration increased beyond 0.02 mM, particularly at 0.1 mM, where the lowest viability was recorded in both 5 and 10% (w/v) GelMA concentrations. This suggests a threshold for cytocompatibility, beyond which reactive oxygen species and radical accumulation may impair cell health. The trends were consistent across both GelMA formulations, though viability appeared slightly increased in the denser 10% GelMA hydrogels, potentially due to their increased stiffness and enhanced mechanical support for cell adhesion and spreading, a phenomenon previously observed in stiffness-sensitive fibroblast cultures. ^40, 41, 69^

LAP is a biocompatible photo-initiator for fabrication of cell-laden GelMA hydrogels. While UV light exposure is generally associated with DNA damage and reactive oxygen species (ROS) generation ^16^; studies have shown that using LAP at lower concentrations, along with minimal UVA irradiation duration, can effectively initiate cross-linking while minimizing cytotoxic effects. ^70^ Here, we observed in 5% (w/v) GelMA samples that cell survival remained relatively stable across concentrations ranging from 0.01 to 0.5% (w/v), with 0.05% (w/v) LAP yielding the highest viability (**Figures 7c**). While a slight downward trend in cell survival was noted with increasing concentrations, none of the conditions exhibited significant cytotoxicity, highlighting LAP’s robustness as a photoinitiator when used with brief UVA exposure (up to 1 minute). Similar trends were observed in the 10% GelMA formulations (**Figure 7d**), where the influence of [LAP] on cell viability was more evident, with significantly higher survival rates. These enhanced viability outcomes in 10% hydrogels can likely be attributed to the increased stiffness of the network, which favors the behavior of stiffness-sensitive fibroblasts.^64^ These results align with previous studies where LAP was shown to maintain high cytocompatibility while enabling efficient and rapid photopolymerization up to 0.5% (w/v) ^46, 70^, making it an attractive choice for a wide range of biomedical applications, including 3D bioprinting and microfabrication. ^24, 31^

The Ru-SPS photoinitiator system is widely recognized for its cytocompatibility, making it a favorable choice for 3D bioprinting and cell-laden hydrogel constructs. Operating within the visible light spectrum (typically around 450 nm), it eliminates the risks associated with UV-induced cellular damage. Although persulfate generates free radicals essential for cross-linking, these radicals are short-lived, and the resulting sulfate ions are considered minimally toxic at the low concentrations typically employed. ^26, 33, 71^ Compared to the LAP system, the Ru/SPS photoinitating system exhibited a narrower biocompatibility window, likely due to increasing [SPS] concentrations with increasing [Ru] (fixed 1:10 molar ratio). Both 5 and 10% (w/v) GelMA formulations (**Figures 7e** and **7f**) maintained high cell viability within the 0.05–0.3 mM range, peaking at 0.25 mM Ru. However, a sharp decline in viability was observed beyond 0.3 mM, indicating a cytotoxic threshold. This sudden drop is presumably attributed to excessive persulfate radical generation, which may disrupt cellular integrity. A previous study has demonstrated that SPS concentrations exceeding 0.3 mM exert significant cytotoxic effects on L929 cells. ^54^ Nevertheless, these same studies also noted that the rapid consumption of SPS during photopolymerization should reduce its residual concentration below toxic levels, even with brief (10-second) light exposure.

We previously reported a plateau effect in the Ru/SPS system: prolonged irradiation beyond a certain threshold did not enhance network stiffness, particularly at Ru concentrations ≥0.5 mM. This interesting phenomenon suggests that the photopolymerization reaction reaches saturation, wherein additional SPS-originated free persulphate radicals can no longer contribute to further cross-linking due to rapid chain termination reactions. As a result, we think that surplus free radicals may accumulate, potentially creating cytotoxicity in cell environment due to unutilized SPS. This hypothesis was further supported by our observations at 1 mM [Ru] with 10 mM [SPS], where both 5% and 10% (w/v) GelMA hydrogels exhibited negligible cell viability, underscoring the cytotoxic effects associated with excessive [SPS] levels. Comparable findings were reported in a separate study that examined the effects of elevated Ru/SPS concentrations (0.5/5 and 1/10 mM) on *ex vivo* lipoaspirate tissues. ^72^ The researchers also mentioned that excessive photoinitiator levels led to excessive free radical generation, resulting in increased chain termination events during cross-linking. This, in turn, caused an accumulation of unreacted free radicals, which negatively impacted tissue viability. Moreover, the study confirmed that SPS concentrations exceeding 0.5 mM exhibited cytotoxic effects on peripheral blood mononuclear cells (PBMCs), further substantiating our observations regarding the cytotoxic threshold of the Ru/SPS photoinitiating system and our related hypothesis.

Overall, increasing GelMA concentration from 5% to 10% appeared to offer marginal improvements in cell viability across all photo-initiator systems, likely due to higher cross-linking density and mechanical stiffness that can influence cell-matrix interactions with fibroblasts. These findings collectively underline the importance of carefully balancing photo-initiator concentration and hydrogel formulation to ensure optimal cytocompatibility. While EY and Ru-SPS systems offer tunability and compatibility with visible light systems, their narrow biocompatibility windows necessitate precise control over formulation conditions. In contrast, LAP provides a broader and more stable platform for photopolymerization, making it especially suitable for applications in regenerative medicine, *in situ* gelation, and cell-laden 3D constructs.

### 2.6 Comparison of Photoinitiation Systems

To comprehensively evaluate the performance of different photoinitiating systems in GelMA hydrogel fabrication, a side-by-side comparison of EY, LAP, and Ru initiators was conducted. Each system presents unique polymerization kinetics, network architectures, and resulting physical properties, which directly influence their suitability for specific biomedical applications. Critical aspects such as mechanical strength, degradation resistance, swelling behavior, and the influence of initiator concentrations were assessed to determine the advantages and limitations of each cross-linking system. **Table 1** below consolidates our findings to provide a clear overview of how each photoinitiator shapes the structure-function relationships in GelMA hydrogels.

**Table 1.** Comparative analysis of GelMA hydrogels crosslinked with EY, LAP, and Ru photoinitiating systems. Optimal photoinitiator concentration ranges were determined based on the findings of this study. Key parameters—including mechanical strength, degradation behavior, swelling capacity, and microstructural features—are summarized across the tested concentration ranges from lowest to highest, highlighting system-specific trends relevant for biomedical applications.

| Parameter | EY-based System | LAP-based System | Ru-based System |
| --- | --- | --- | --- |
| Light Source | Visible light (green, 450-550 nm) | UV light (365 nm) or Visible light (405 nm) | Visible light (blue, 400-450 nm) |
| Optimal Concentration Range | 0.005 to 0.1 mM | 0.01 to 0.5% (w/v) | 0.05 to 0.5 mM |
| Polymerization Kinetics | Moderate (<40s), efficient and homogeneous cross-linking | Fast(<10s), highly-efficient but heterogeneous cross-linking | Fast (<10s), less-efficient, heterogeneous cross-linking |
| Mechanical Strength (kPa) | ~4 to 10 kPa : (5% GelMA)<br>~25 to 45 kPa : (10% GelMA) | ~0.5 to 3.2 kPa : (5% GelMA)<br>~8 to 30 kPa : (10% GelMA) | ~0.2 to 2.4 kPa : (5% GelMA)<br>~1 to 17.5 kPa : (10% GelMA) |
| Swelling Ratio (%) (5% GelMA) | Lowest (~550–1000%) : (5% GelMA)<br>Lowest (~375–450%) : (10% GelMA) | High (~1000–1700%) : (5% GelMA)<br>Moderate (~550–1200%) : (10% GelMA) | Highest (~1100–1900%) : (5% GelMA)<br>Moderate (~600–1100%) : (10% GelMA) |
| Degradation Behavior (h) ( 5% GelMA) | Controlled, slow degradation (105-195 min) : (5% GelMA)<br>Controlled, slow degradation (195-360 min) : (10% GelMA) | Controlled, moderate degradation (60-135 min) : (5% GelMA)<br>Controlled, slow degradation (195-360 min) : (10% GelMA) | Uncontrolled, rapid degradation (45-90 min) : (5% GelMA)<br>Controlled, moderate degradation (90-255 min) : (10% GelMA) |
| Pore Morphology (SEM) | Small, uniform, highly interconnected pores | Large, thick, irregular pores; some heterogeneity | Thinner walls, some collapse, irregular connectivity |
| Cytocompatibility (%) (w/ NIH-3T3 cells) | High (Visible light, mild photoinitiator system) | High, but UV exposure may be limiting for sensitive cells | High (Visible light, toxic at elevated [SPS]) |
| Hydrogel Network Architecture | Well-distributed, balanced structure | Dense, locally crosslinked, variable homogeneity | Loosely connected, lower crosslink density |
| Advantages of the System | Easily tunable stiffness profile<br>Visible-light, more cytocompatible *in vivo*<br>Comparatively high stiffness, slower degradation | Fast gelation profile<br>Only one parameter, easier to modify<br>High water solubility | Fast gelation profile<br>Visible-light, more cytocompatible *in vivo*<br>Favorable environment for many sensitive cell types |
| Disadvantages of the System | Requires co-initiator and comonomer, complex system<br>Cytotoxic at elevated TEA or NVP concentrations<br>pH-sensitive photo-crosslinking mechanism | Cytotoxic at prolonged UV exposure<br>Limited tunability<br>Uneven cross-linking and heterogeneous network | Complex handling, due to sensitivity to oxygen<br>Lower mechanical strength of hydrogels<br>Potential cytotoxicity at elevated [SPS] |
| Application Suitability | Cell-laden scaffolds<br>Drug-delivery hydrogel systems<br>Wound healing | 3D bioprinting<br>Microfluidic lab-on-a-chips<br>Injectable hydrogel systems | Cell encapsulations, bioprinting<br>Cardiac & ocular tissue engineering<br>Injectable, rapidly degradable systems |

## 3. Conclusion

This study provides the first comprehensive comparative analysis of three prominent photoinitiator systems (EY, LAP, and Ru), on the mechanical, degradation, swelling, and cytocompatibility profiles of GelMA hydrogels. By systematically varying photoinitiator concentrations and evaluating their effects on both 5% and 10% (w/v) GelMA formulations, we present critical insights into how photoinitiator chemistry, dosage, and crosslinking dynamics synergistically influence hydrogel performance and ultimately dictate their biomedical applicability.

Each photoinitiator system demonstrated unique behavior in a concentration-dependent manner, reflecting the complexity of their underlying photopolymerization mechanisms. The EY/TEA/NVP system, operating through a Type II photoinitiation mechanism under green light, exhibited an optimal stiffness at 0.02 mM EY, beyond which both mechanical integrity and cell viability declined. This decline is attributed to radical overproduction leading to network heterogeneity, premature chain terminations, and possible triplet-triplet annihilation of EY molecules. Nevertheless, the EY system facilitated the formation of homogenous and highly crosslinked networks with superior enzymatic resistance and low swelling, suggesting its high suitability for cell-laden scaffolds, drug delivery depots, and wound healing patches, where longer-term mechanical stability and structural integrity are critical.

In contrast, the LAP system, which operates through a rapid Type I photoinitiation process upon UV light exposure, showed atypical mechanical behavior at low concentrations (0.01–0.02% w/v), outperforming higher concentrations in storage modulus over longer irradiation times. This surprising outcome suggests a more efficient and homogeneous network formation under slower radical generation, while higher concentrations exhibited diminished mechanical properties due to rapid and excessive radical generation, leading to chain termination and dangling polymer ends. Still, LAP’s wide biocompatibility range, fast gelation, and high resolution under short UV exposure reinforce its robustness as a photoinitiator. These attributes render LAP-based GelMA hydrogels ideal candidates for 3D bioprinting, microfluidic applications, and injectable hydrogel systems that demand rapid setting, spatial control, and moderate mechanical properties.

The Ru/SPS system, leveraging a photoredox mechanism under blue light, emerged as a promising yet delicate platform, with stiffness increasing up to 0.5 mM Ru, after which mechanical properties plateaued or even deteriorated due to saturation in radical propagation. The fixed Ru:SPS molar ratio (1:10) proved critical for ensuring balance between network density and cytocompatibility. Excessive SPS levels at higher Ru concentrations caused radical accumulation and cytotoxicity, emphasizing the importance of precise stoichiometric control. Ru-crosslinked hydrogels displayed the highest swelling and fastest enzymatic degradation, indicative of looser, heterogeneous networks. These features, combined with the system’s favorable cytocompatibility at lower concentrations, highlight its potential for bioprinting of cell-laden constructs, cardiac and ocular tissue engineering, and rapidly-degradable injectable systems where transient scaffold support is desired.

Importantly, the results also demonstrate that GelMA concentration modulates photoinitiator behavior, as 10% (w/v) hydrogels consistently exhibited higher stiffness, lower swelling, and enhanced degradation resistance, attributed to higher methacryloyl group density. Across all systems, increased GelMA content also modestly improved cell viability, particularly relevant for applications using fibroblast-sensitive matrices.

Taken together, this study underlines that photoinitiator type and concentration are not interchangeable parameters but must be specifically optimized depending on the intended biomedical application. Our findings offer an invaluable reference for researchers seeking to tailor GelMA hydrogels by rational photoinitiator selection. Moreover, the correlation between stiffness, degradation, and swelling revealed in this study provides a mechanistic understanding for designing hydrogels that meet the demands of structurally complex, biologically responsive, and clinically translatable tissue engineering platforms.

Future studies may further benefit from integrating photo-polymerization kinetics models and machine learning tools to predict and optimize hydrogel behavior under various photoinitiating conditions. Expanding the scope of visible-light photoinitiators, particularly those with minimal cytotoxic byproducts, also holds promise for advancing GelMA-based hydrogel systems toward clinical and translational readiness.

## 4. Experimental Section

### 4.1. Materials

All materials used in this study have been obtained from Sigma Aldrich unless otherwise mentioned.

### 4.2. Preparation of GelMA Macromonomers

GelMA (Gelatin Methacryloyl) was synthesized according to protocol, by dissolving bovine skin gelatin (Type B, ∼225g bloom) in a carbonate-bicarbonate (CB) buffer solution at 50°C. ^73^ Methacrylic anhydride (MA) was then added dropwise to the gelatin solution while stirring, and the mixture was maintained at 50°C for 3 hours. After adjusting the pH to 7.4, the solution was dialyzed against distilled water for 7 days at 40°C to remove any unreacted MA. The dialyzed solution was then lyophilized to obtain a dry powder form of GelMA, which was stored at -20°C for future use.

### 4.3. GelMA Characterization

*H-NMR Analysis.* H-nuclear magnetic resonance spectrometer (^1^H-NMR) was preferred to determine the degree of methacrylation in synthesized GelMA. For this purpose, 10 mg of both gelatin and GelMA were dissolved in 0.6 mL of deuterium oxide (D_2_O) at 25 °C, separately and then analyzed by a Bruker NMR 500 MHz Advance spectrometer (Bruker BioSpin Rheinstetten, Germany). The degree of gelatin modification with methacrylamide is influenced by the availability of amino functional groups within the gelatin molecule, both before and after the modification process. This suggests a direct correlation between the extent of modification and the number of reactive amino groups susceptible to methacrylamide attachment. Thus, to normalize the amine signals (2.9 ppm) of methacrylated arginine, the phenylalanine signals (7.0-7.5 ppm) were used as an internal reference. This internal reference remained unaffected by the reaction with methacrylic anhydride across all samples. The degree of methacrylation was subsequently calculated by normalizing the amine signal at 2.9 ppm to the phenylalanine signals at 7.0-7.5 ppm, using the following equation:

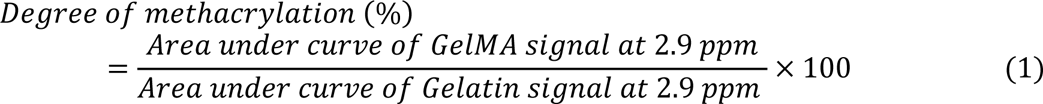

#### FTIR Analysis

Fourier-transform infrared spectroscopy (FTIR, Nicolet iS10 instrument, ThermoFisher, United States) was employed to analyze the chemical composition of gelatin and synthesized GelMA with a resolution of 4 cm^-1^. By examining the absorption peaks and frequencies of average of 64 scans in the infrared spectra, covering a range of 600 to 4000 cm^-1^, the specific molecular groups present within these biomaterials were identified. In short, FTIR spectroscopy was utilized to verify the successful incorporation of methacrylate groups into the GelMA polymer.

#### XRD Analysis

X-ray diffraction (XRD) analysis was employed to investigate the crystalline or amorphous nature of GelMA sample before and after methacrylation. XRD patterns were acquired using a Bruker D2 Phaser X-Ray Diffractometer equipped with graphite-filtered Cu Kα radiation (λ = 1.5418 Å) operating at 40 kV and 20 mA. Data was collected over a 2θ range of 5° to 80° with a step size of 0.021° and a scan rate of 0.8 s per step.

#### Morphological Analysis

Size, porosity, and surface morphology of the GelMA and selected GelMA hydrogels were visualized using a ZEISS Ultra Plus Field Emission Scanning Electron Microscope (FE-SEM, ZEISS, Oberkochen, Germany). Before SEM analysis, GelMA hydrogels were lyophilized overnight and cut into thin slices. Additionally, all the samples were gold-sprayed before SEM scanning.

### 4.4. GelMA Prepolymer Solution Preparation

All prepolymer solutions were freshly prepared on the day of the experiment and maintained at 37°C in dark conditions to preserve stability. To prepare 5% (w/v) GelMA solutions with varying EY concentrations (0.005, 0.01, 0.02, 0.04, 0.05, and 0.1 mM), 50 mg of lyophilized GelMA was first dissolved in calculated volume of PBS (pH 7.4). Then, 100 µL of 1 M TEA, 50 µL of 1 M NVP and appropriate volume of EY stock solution were subsequently added to adjust the final volume to 1 mL (resulting in final TEA and NVP concentrations of 100 mM and 50 mM). For the preparation of LAP-based 5% (w/v) GelMA prepolymer solutions, 50 mg of lyophilized GelMA was initially dissolved in appropriate volume of PBS at 37°C. Subsequently, precisely calculated volumes of LAP stock solution were added to achieve final LAP concentrations (0.01, 0.02, 0.05, 0.07, 0.1, 0.15, 0.25, 0.5 % (w/v)) and to adjust the total volume of prepolymer solution to 1 mL. Lastly, to prepare 5% (w/v) GelMA prepolymer solutions with changing Ru concentrations (0.02, 0.05, 0.1, 0.2, 0.25, 0.3, 0.5, 1 mM), 50 mg of lyophilized GelMA was first dissolved in predetermined volume of PBS at 37°C. Subsequently, calculated volume of 50 mM SPS stock solution was added to maintain a 1:10 molar ratio between Ru and SPS, followed by the addition of appropriate volume of Ru stock solution to adjust the total volume of prepolymer solution to 1 mL.

### 4.5. Rheological Properties of Hydrogels

All GelMA prepolymer samples were freshly prepared on the day of the experiment and incubated at 37°C. The stiffness of GelMA prepolymer solutions was measured at 25 °C using a Discovery HR-2 Rheometer (TA Instruments, Delaware, USA) equipped with parallel-plate geometry (20 mm in diameter) and a gap size of 1 mm. To induce *in situ* cross-linking of 5 and 10% (w/v) GelMA prepolymer solutions, 3 different light-induced systems, which utilizes LAP, EY, and Ru as photoinitiators, were applied via custom-built LED attachments of varying wavelength of light (TA Instrument, 365nm, 430nm, 534 nm, 10 mW/cm^2^, respectively). Before rheological analysis, to make sure that each changing light source provides equal amount of light intensity (10 mW/cm^2^), each light sources’ corresponding ampere value was optimized through power measurements via PM100D power meter (THORLABS, New Jersey, USA) (**Figure S5**). The cross-linking process was monitored through a time sweep oscillatory test performed under a constant strain amplitude and frequency within the linear viscoelastic region (LVER), and triplicate measurements were recorded. The optimal strain percentage for 5% and 10% (w/v) GelMA prepolymer solutions within LVER region was evaluated using an oscillatory strain sweep test, applying strain values ranging from 1 to 100% and 1 to 200% at a constant frequency of 1 Hz. Similarly, the optimal frequency range for the viscoelastic analysis of each photoinitiating system’s selected main conditions was determined through a frequency sweep test, conducted at a constant strain of 0.01% (**Figure S6**). Based on our analysis, we determined the optimal strain and frequency for all rheological measurements of 5% and 10% GelMA hydrogels to be 0.5% and 5 Hz, respectively.

A sample volume of 200 µL was loaded onto the attachment for the test. For Young’s modulus calculations, following Equation 2 was used:

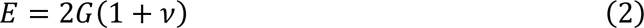

where E, G, and *ν* represent the Young’s modulus, storage modulus, and Poisson’s ratio, respectively. For the GelMA hydrogel, Poisson’s ratio was taken as 0.5; thus, E is approximated to 3G. ^74^

### 4.6. Preparation of Hydrogels via Light-induced Cross-linking

5 and 10% (w/v) GelMA hydrogels were prepared similarly for each condition of three different photoinitiator systems. Basically, arranged prepolymer solutions were pipetted to a 96-well plate in triplicates and cross-linked using our handmade LED light sources, irradiating 365, 430, and 532 nm of light for LAP, Ru, and EY-based hydrogel photopolymerization, respectively (constant 10 mW/cm^2^). While EY-based hydrogels were crosslinked for 5-minutes to ensure completed polymerization of GelMA, LAP and Ru hydrogels were obtained by 1-minute cross-linking in 96-well plate.

### 4.7. *In vitro* Enzymatic Degradation Study

To assess *in vitro* biodegradability behavior of the hydrogels, the preweighted hydrogel samples (W_0_) were immersed in solution containing 0.1% Collagenase from *Clostridium histolyticu* (C6885) and incubated at 37 °C. At each time point (15, 30, 45, 60, 75, 90, 135, 195, 255, 360 minutes), produced GelMA hydrogels were carefully taken out from the solution, blot-dried with Kimtech wipe, and their weight (W_F_) was measured. Equation 3 was applied to calculate the gel retention (n = 3) of the hydrogel samples:

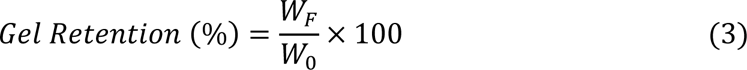

where W_F_ refers to the weight of hydrogel sample at specific time point and W_0_ represents the weight of hydrogels before immersing the samples into the enzyme solution.

### 4.8. Swelling Study

The swelling behavior of GelMA hydrogels was determined by immersing the samples in PBS solutions and incubating them at 37 °C. After the preparation, hydrogels were frozen and lyophilized to obtain dry weight of the hydrogels (W_0_). Subsequently after dry weight measurements, hydrogels were allowed to swell in PBS solutions at 37°C. The swollen hydrogel samples were withdrawn, blot-dried with Kimtech wipe, and weighted at predetermined time points (1, 2, 3, 4, and 24 h). All measurements were completed in triplicates. The following Equation 4 was utilized to calculate the swelling ratio of hydrogel samples (n = 3):

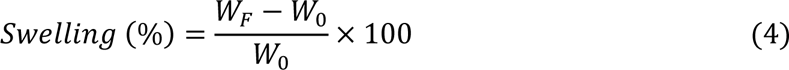

where W_F_ is the weight of the swollen hydrogel at time points, while W_0_ demonstrates the weight of the freeze-dried hydrogel sample before immersing it into the PBS solution.

### 4.9. Cell Culture

NIH-3T3 fibroblast cells were cultured in DMEM High Glucose medium, supplemented with 10% fetal bovine serum (FBS) and 1% Penicillin-Streptomycin (Pen-Strep), and maintained at 37°C in a 5% CO_2_ incubator. Once the cells reached confluence, they were collected by incubating with 0.25% trypsin-EDTA for 5-minutes, followed by centrifugation at 200 × g for 7 minutes to collect the cell pellet.

### 4.10. Cell Cytotoxicity

To evaluate the effects of GelMA hydrogels crosslinked with different EY, LAP, and Ru concentrations on fibroblast cell viability, a Cell Titer-Glo (CTG) assay (Promega) was performed on day 7, following the manufacturer’s instructions. Fibroblasts were initially seeded at a density of 12000 cells per well in a 96-well plate and allowed to adhere. On day 7, cell containing GelMA hydrogels were gently washed once with PBS, and CTG reagent was subsequently added to each well. After allowing the luminescent signal to stabilize for 10 minutes, the signal was measured using a microplate reader to assess cell viability.

### 4.11. Statistical Analysis

A one-way ANOVA with Tukey comparison test on GraphPad Prism Software (Graph-Pad Software version 10.4.1, CA, USA) was preferred for statistical significance analysis. A value of p ≤ 0.05 was considered as statistically significant for resulting data evaluation. The significance levels are denoted as follows: n.s. for p > 0.05, * for p ≤ 0.05, ** for p ≤ 0.01, *** for p ≤ 0.001, and **** for p ≤ 0.0001

## Conflict of Interest

The authors declare no conflict of interest.

## Data Availability Statement

The data supporting this study’s findings are available from the corresponding author upon reasonable request.

## Supporting Information

The Supporting Information is available free of charge.

## CRediT authorship contribution statement

**D.D:** Writing – original draft, Methodology, Investigation, Data curation, Visualization Conceptualization. **I.C.K:** Writing – original draft, Methodology, Conceptualization, Validation. **S.K*:** Writing – review & editing, Writing – original draft, Supervision, Resources, Project administration, Funding acquisition, Conceptualization.

## Acknowledgement

The authors sincerely acknowledge the support and use of the facilities and services provided by the Koç University Research Center for Surface Science (KUYTAM) and the Koç University Research Center for Translational Medicine (KUTTAM). This research was financially supported by the Scientific and Technological Research Council of Türkiye (TÜBİTAK) **Support.** Additionally, D.D. and I.C.K. gratefully acknowledge the BIDEB scholarships awarded by TÜBİTAK.

## Notes

### Competing Interest Statement

The authors have declared no competing interest.

