## Supplementary information for "Effect of Photoinitiation Process on Photo-Crosslinking of Gelatin Methacryloyl Hydrogel Networks"

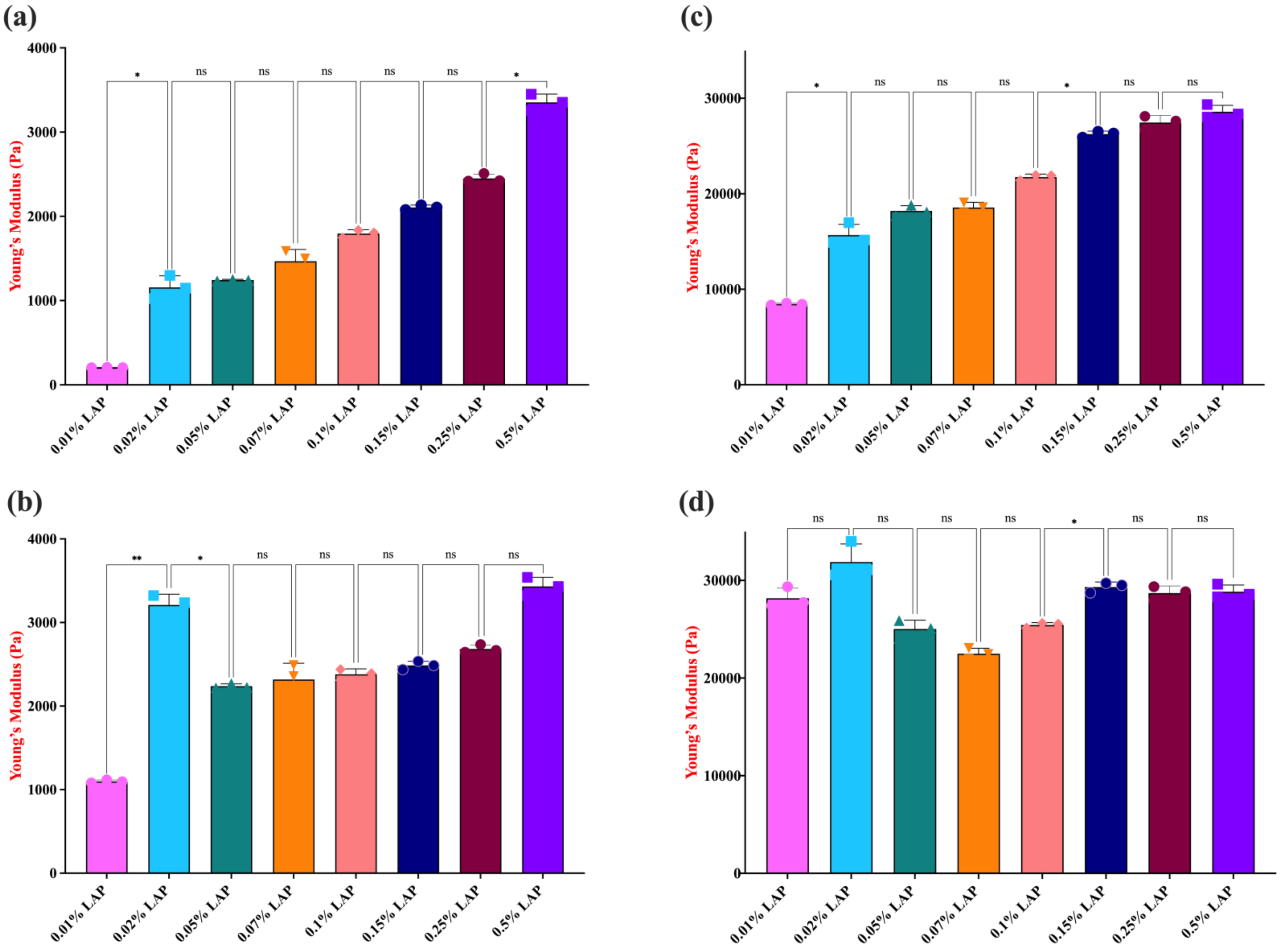


**Figure S1**. Influence of changing UV light exposure duration (1-minute vs. 5-minute) on stiffness of LAP-based hydrogels. Young's moduli values for 5% (w/v) GelMA hydrogels cross-linked for 1- and 5-minute (**a, b**) and 10% (w/v) GelMA hydrogels cross-linked for 1- and 5-minute (**c, d**) were calculated through the independent replicates of storage modulus (*G'*) data. Statistical analysis was conducted using a one-way ANOVA followed by Tukey's post hoc test, with a significance threshold of $p \leq 0.05$. Statistical notations are as follows: n.s. (not significant, $p > 0.05$), * ($p \leq 0.05$), and **** ($p \leq 0.0001$). Error bars shown (n=3) indicate the standard deviation.

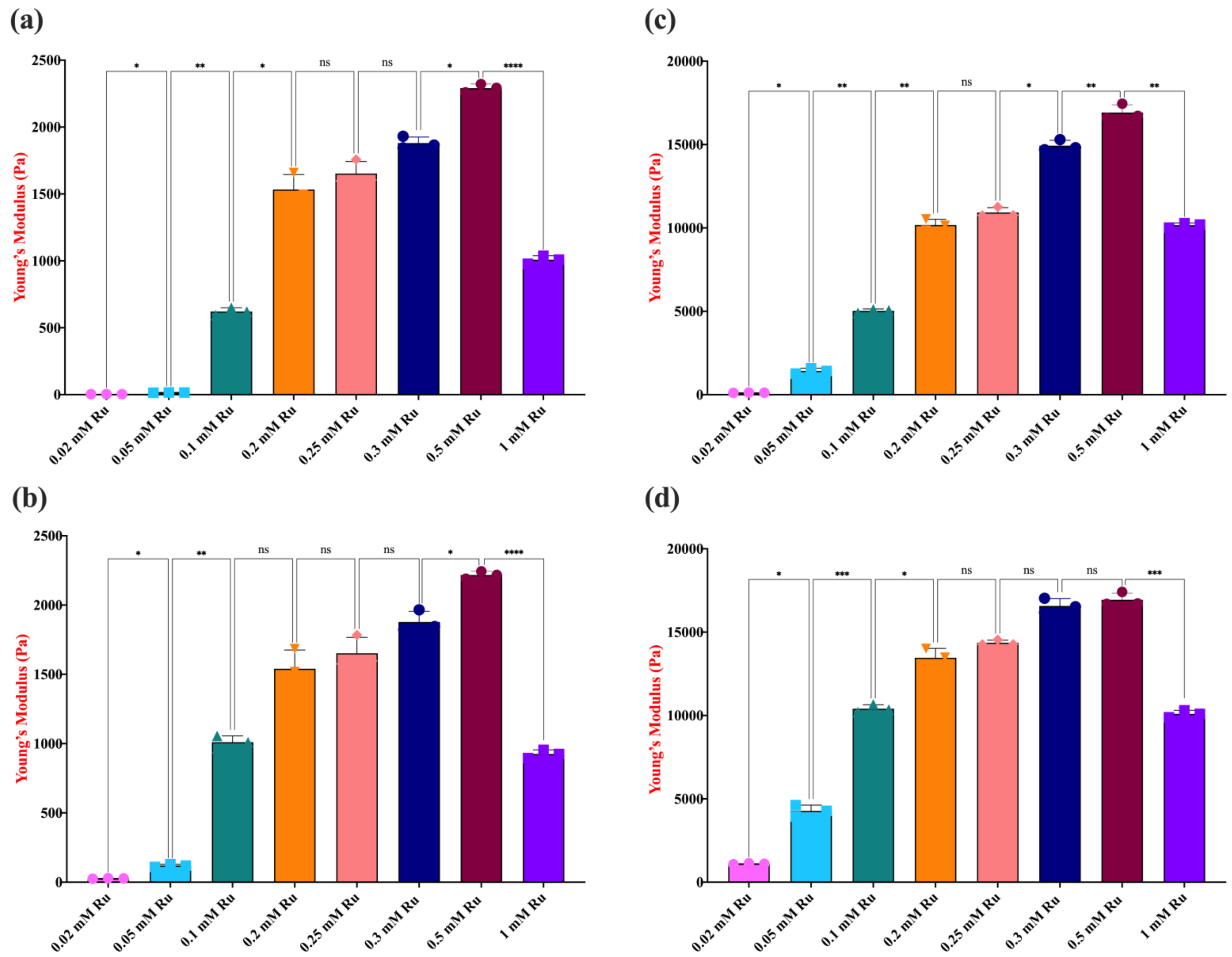


**Figure S2.** Influence of changing blue light ($\lambda$= 430 nm) exposure duration (1-minute vs. 5-minute) on stiffness of Ru-based hydrogels. Young's moduli values for 5% (w/v) GelMA hydrogels cross-linked for 1 and 5 minutes (**a, b**) and 10% (w/v) GelMA hydrogels cross-linked for 1 and 5 minutes (**c, d**) were calculated through the independent replicates of storage modulus (*G'*) data. Statistical analysis was conducted using a one-way ANOVA followed by Tukey's post hoc test, with a significance threshold of $p \leq 0.05$. Statistical notations are as follows: n.s. (not significant, $p > 0.05$), * ($p \leq 0.05$), and **** ($p \leq 0.0001$). Error bars shown (n=3) indicate the standard deviation.

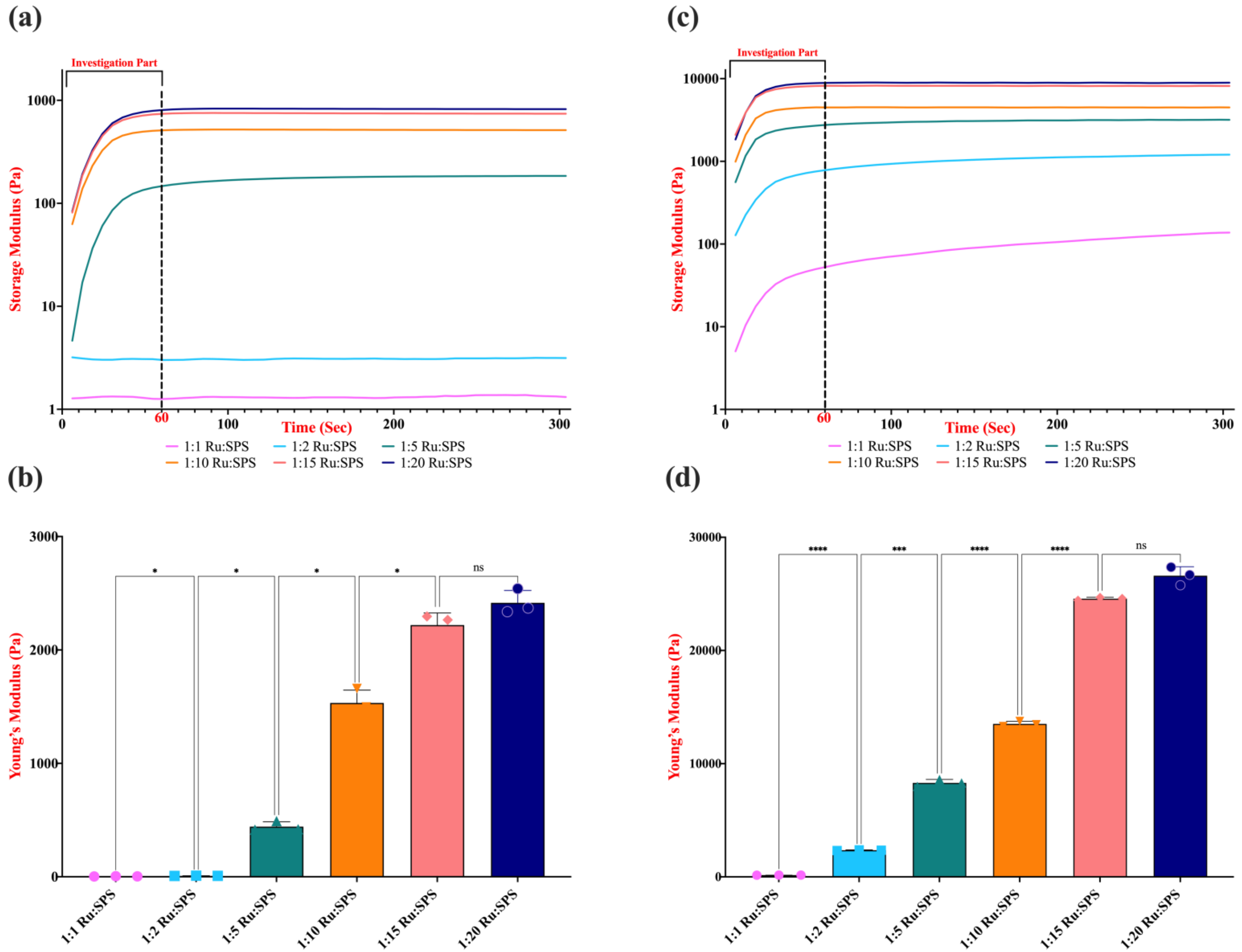


**Figure S3.** Influence of changing [Ru]:[SPS] on 5 and 10% (w/v) GelMA hydrogels by keeping [Ru] constant as 0.2 and 0.3 mM, respectively. Hydrogels were obtained through photopolymerization on a parallel plate rheometer at room temperature under blue light conditions ($\lambda$= 430 nm for Ru), at an intensity of 10 mW/cm$^2$ for 5 minute to measure real-time Storage modulus and Young's modulus of 5% (**a, b**) and 10% (w/v) GelMA hydrogels (**c, d**). The data represent the average and smoothed storage modulus ($G'$) obtained from three independent replicates. Statistical analysis was conducted using a one-way ANOVA followed by Tukey's post hoc test, with a significance threshold of $p \leq 0.05$. Statistical notations are as follows: n.s. (not significant, $p > 0.05$), * ($p \leq 0.05$), and **** ($p \leq 0.0001$). Error bars shown (n=3) indicate the standard deviation.

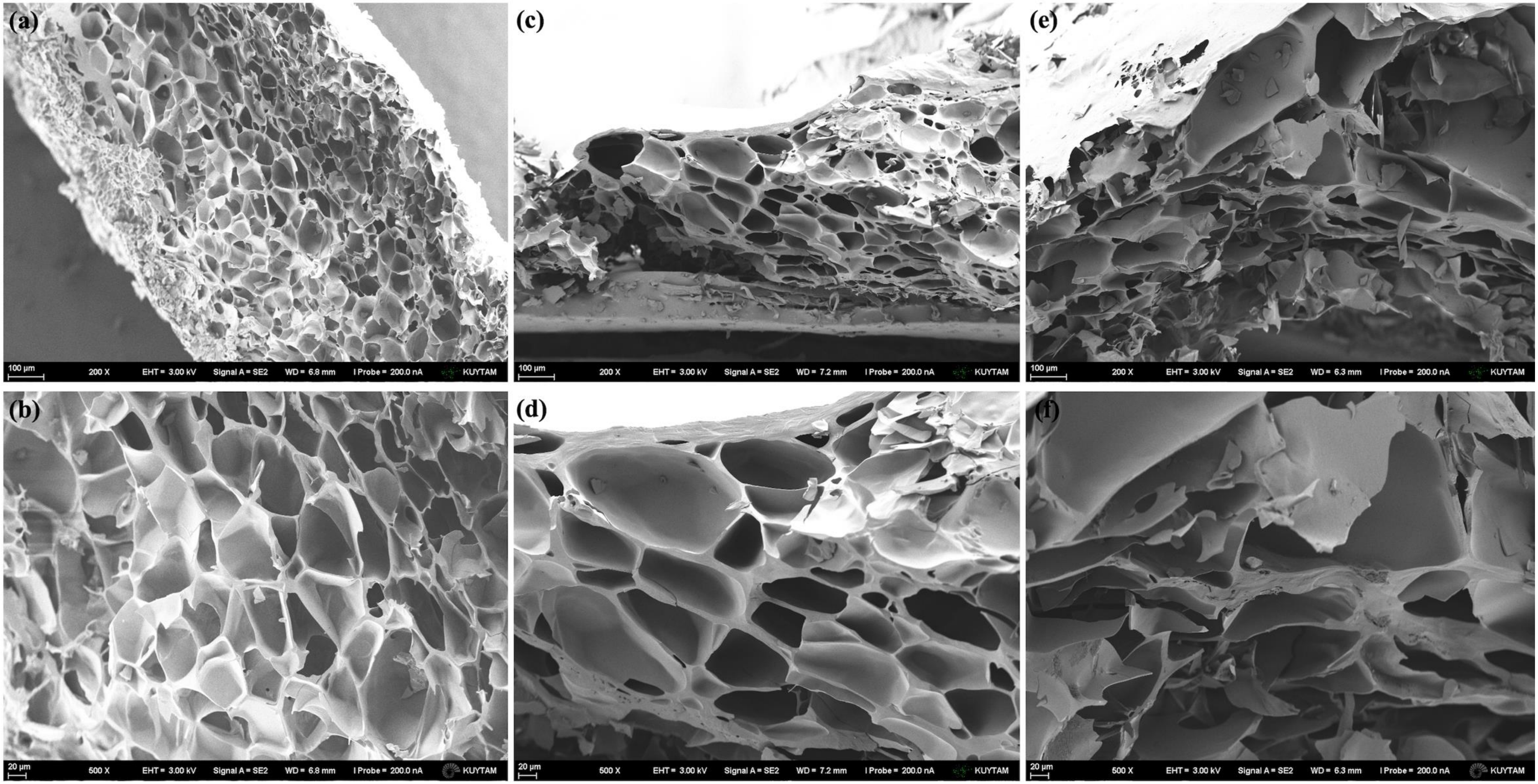


**Figure S4**. FE-SEM imaging was conducted to examine the microstructural characteristics of 5% (w/v) GelMA hydrogels polymerized using different photoinitiating systems. Hydrogels were prepared with 0.01 mM EY- (**a, b**), 0.15% wt. LAP- (**c, d**), and 0.3 mM Ru-based photo-crosslinking (**e, f**) and analyzed at magnifications of 200× and 500×. These specific conditions were selected as they represent intermediate concentrations within each photoinitiator system while exhibiting comparable Young's moduli, allowing for a direct comparison of the morphological differences induced by distinct polymerization mechanisms.

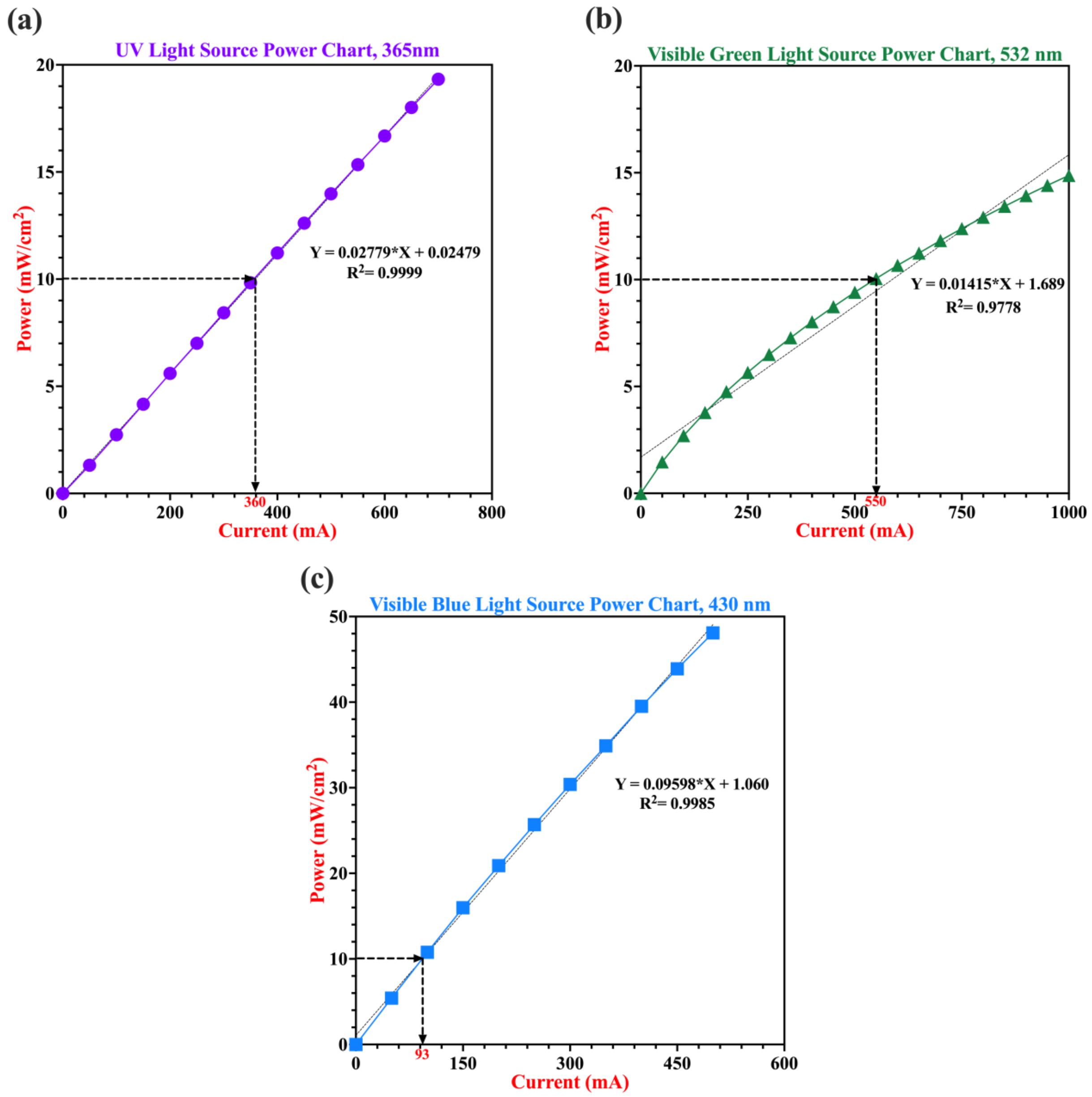


**Figure S5**. Calibration of hand-made light sources based on power–current (ampere) relationships. Power output (mW/cm$^2$) was measured as a function of current (mA) for each light source using a power meter to ensure consistent light intensity delivery. Target intensity of 10 mW/cm² was achieved at 360 mA for 365 nm UV light (**a**), 550 mA for 532 nm visible green light (**b**), and 93 mA for 430 nm visible blue light (**c**). All measurements were performed at a constant distance (~2 cm) between the light source and the power meter probe and were routinely verified throughout the study to maintain experimental consistency.

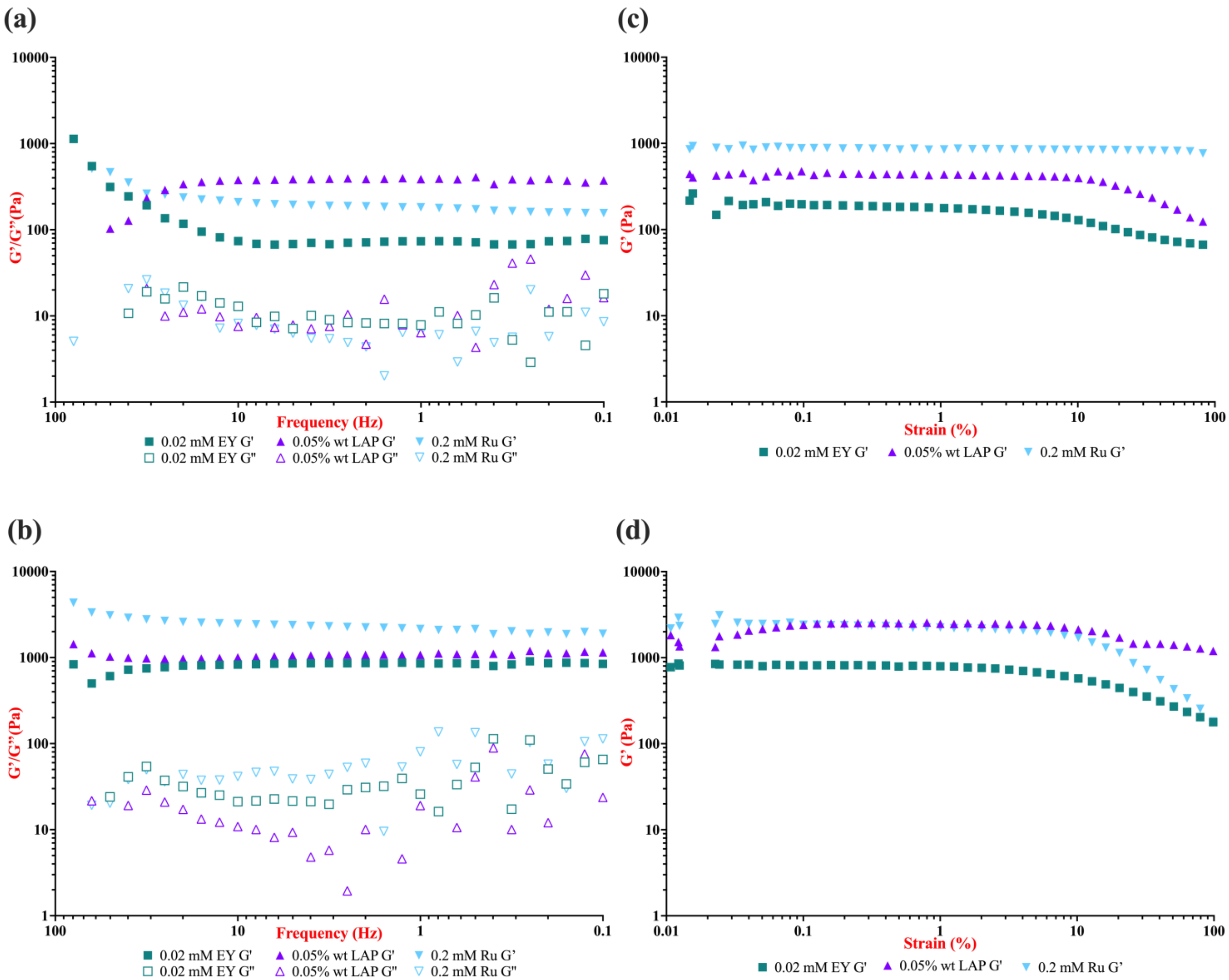


**Figure S6**. Viscoelastic properties of 5 and 10% (w/v) GelMA hydrogels at selected conditions of EY, LAP and Ru-photoinitiating system. (**a, b**) Frequency sweep analysis of 5% and 10% (w/v) GelMA hydrogels polymerized with 0.02 mM EY, 0.05% wt. LAP, and 0.2 mM Ru under light exposure at 530 nm, 365 nm, and 430 nm, respectively, with an intensity of 10 mW/cm². The data illustrate the evolution of storage modulus (*G′*) and loss modulus (*G″*) as a function of frequency (0.1 to 100 Hz). (**c, d**) Strain sweep analysis of 5% and 10% (w/v) GelMA hydrogels prepared under the same conditions to determine the linear viscoelastic (LVE) region. The data represent the corresponding storage modulus (*G′*) as a function of strain %, ranging from 0.01% to 100% for 5% GelMA and from 0.01% to 200% for 10% GelMA.
